# GranatumX: A Community-engaging, Modularized, and Flexible Webtool for Single-cell Data Analysis

**DOI:** 10.1101/385591

**Authors:** David Garmire, Xun Zhu, Aravind Mantravadi, Qianhui Huang, Breck Yunits, Yu Liu, Thomas Wolfgruber, Olivier Poirion, Tianying Zhao, Cédric Arisdakessian, Stefan Stanojevic, Lana X. Garmire

**Affiliations:** Department of Electrical & Computer Engineering, University of Michigan, Ann Arbor, MI48105, USA; Epidemiology Program, University of Hawaii Cancer Center, Honolulu, HI 96813, USA; Department of Computational Medicine and Bioinformatics, University of Michigan, Ann Arbor, MI48105, USA; Department of Biostatistics, University of Michigan, Ann Arbor, MI48109, USA; Molecular Biosciences and Bioengineering Graduate Program, University of Hawaii at Manoa, Honolulu, HI 96822, USA

**Keywords:** single cell, RNA-Seq, analysis, software, pipeline, plugin, webtool

## Abstract

We present GranatumX, a next-generation software environment for single-cell data analysis. GranatumX is inspired by the interactive web tool Granatum. It enables biologists to access the latest single-cell bioinformatics methods in a web-based graphical environment. It also offers software developers the opportunity to rapidly promote their own tools with others in customizable pipelines. The architecture of GranatumX allows for easy inclusion of plugin modules, named Gboxes, that wrap around bioinformatics tools written in various programming languages and on various platforms. GranatumX can be run on the cloud or private servers and generate reproducible results. It is a community-engaging, flexible, and evolving software ecosystem for scRNA-Seq analysis, connecting developers with bench scientists. GranatumX is freely accessible at http://garmiregroup.org/granatumx/app.

## Introduction

Single-cell RNA sequencing (scRNA-seq) technologies have advanced our understanding of cell-level biology significantly [1]. Many exciting scientific discoveries are attributed to new experimental technologies and sophisticated computational methods [2,3]. Despite the progress in cultivating professionals with cross-discipline training, a gap continues to exist between the wet-lab biology and the bioinformatics community. Moreover, with the rapid development of many varieties of modules handling different parts of scRNA-seq analysis [4–6], it becomes increasingly challenging for bioinformaticians themselves to decide which method to choose. Although some analytical packages such as SINCERA [7], Seurat [8], and Scanpy [9] provide complete scRNA-seq pipelines, they require users to be familiar with their corresponding programming language (typically R or Python), installation platform, and command-line interface. This overhead hinders wide adoption by experimental biologists, especially those newly adopting scRNA-seq technologies. A few platforms, such as ASAP [10] and our own previous tool Granatum [11], provide intuitive graphical user interfaces and may be useful for a first-hand exploratory check. However, Granatum does not allow for modularity, while ASAP lacks flexibility and restricts the user to a set number of computational tools. Here we present GranatumX, the new generation of scRNA-seq analysis platforms that aims to solve these issues systematically. Its architecture facilitates the rapid incorporation of cutting-edge tools and enables the efficient handling of large datasets aided by virtualization [12].

## Methods

### Architectural overview

GranatumX consists of three independent components: Central Data Storage (CDS), User Interface (UI), and Task Runner (TR). CDS stores all data and metadata in GranatumX, including the uploaded files, processed intermediate data, and final results. The other two components of GranatumX both have controlled access to the central data storage, which allows them to communicate with each other. CDS is implemented using a PostgreSQL database and a secure file system based data warehouse. UI is the component with which wet-lab biologists interact. The layout is intuitive with Gbox settings while providing a flexible and customizable analysis pipeline. UI also allows for the asynchronous submission of tasks before they can be run by the back-end. UI is implemented using JavaScript, with the ReactJS framework. The submitted jobs queue up in the database and can be retrieved in real-time by TR. TR monitors the task queue in the CDS in real-time, actively retrieves the high-priority tasks (based on submission time), initializes the corresponding Gboxes, and prepares the input data by retrieving relevant data from CDS.

### Deployment

GranatumX uses Docker to ensure that all Gboxes can be installed reproducibly with all their dependencies. As a result, GranatumX can be deployed in various environments including personal computers, dedicated servers, High-Performance Computing (HPC) platforms, and cloud services. The installation instructions are detailed in the README file of the source code.

### Responsive UI

The web-based UI offers different device-specific layouts to suit a wider range of screen sizes. On Desktop computers, the UI takes advantage of the screen space and uses a panel-based layout, and maximizes the on-screen information. On small tablets and mobile devices with limited screen space, a collapsible sidebar-based layout is used to allow the most important information (the results of the current step) to show up on the screen.

### Recipe system

Most studies can use similarly structured pipelines, which typically consist of data entry (upload and parsing), data pre-processing (imputation, filtering, normalization, etc.), and finally data analysis functionalities (clustering, differential expression, pseudo-time construction, network analysis, etc.). GranatumX allows users to save a given pipeline into a “recipe” for the future. GranatumX comes with a set of built-in recipes, which cover many of the most common experiment pipelines.

### Software Development Kits (SDKs)

GranaumX SDKs are made for Python and R. These SDKs provide a set of Application Programming Interfaces (APIs) and helper functions that connect Gbox developer’s own code with the core of GranatumX The detailed documentation can be found in the Github repository.

There are three steps to build a new Gbox from the existing code: 1) Write an entry point in the language of the developer’s choice. The entry point uses the SDK to retrieve necessary input from the core of GranatumX and send back output to the core after the results are computed. 2) Package the entry point, the original package source code, and any dependencies into a docker image using a Dockerfile and the “docker build” command. 3) Write a UI specification for the Gbox. The specification is a simple YAML file that declares the data requirements of the Gbox.

### Pipeline customization

GranatumX allows for full customization of the analysis pipeline. An analysis pipeline has a number of Gboxes organized in a series of steps. Note that two different steps can have the same underlying Gbox. For example, two PCA Gboxes can appear before and after imputation, to evaluate its effect. Because the data are usually processed in a streamlined fashion, later steps in the pipeline usually depend on data generated by the earlier steps. Steps can be added from the app-store into the current project and can be removed from the pipeline at any time. A newly added step can be inserted at any point in the pipeline and can be reordered in any way, as long as such re-arrangement does not violate the dependency relationships.

### Current GranatumX cloud server setup

The current GranatumX web version is hosted on OVHcloud, with specs: Intel Haswell vCPU 128 GB RAM Xeon E5-1650 4GHz. Additionally the https protocol is verified with Let’s Encrypt (https://letsencrypt.org) with an Apache 2 server (https://httpd.apache.org/) and a site registered with No-IP (https://www.noip.com/). This server uses a proxy implementation to pass a user to the Node.js web service. In this manner, Node does not have to manage the security or https connections which allows setup to occur efficiently in an enterprise system. Additionally, an optional fast compute system may be connected to the OVH cloud server through ssh tunneling which allows the local port to be mapped to the remote connection. In this manner, a high speed rig can be connected -- in this case, the AMD 3590x can be connected without having to procure a new cloud system.

### Project management

The studies in GranatumX are organized as projects. Each user can manage multiple concurrent projects. The automatic customer’s report can be generated per project using the parameters and results stored in the CDS.

### Example data sets

Three datasets are used in this report. One data set is downloaded from GSE117988, a study on a patient with metastatic Merkel cell carcinoma, treated using T cell immunotherapy as well as immune-checkpoint inhibitors (anti-PD1 and anti-CTLA4) but later developed resistance [15]. A second dataset is Tabula Muris dataset, which contains 54,865 cells from 20 organs and tissues of the mouse [16]. Another dataset is the 1.3 Million Brain Cells from E18 Mice, downloaded from 10x genomics website: https://support.10xgenomics.com/single-cell-gene-expression/datasets/1.3.0/1M_neurons (accessed on date 05/09/2020). This dataset contains 1,308,421 cells from embryonic mice brains, done by Chromium™ Single Cell 3’ Solution (v2 Chemistry).

### GranatumX Plug-in development

The detailed instruction document, the tutorial youtube or youku videos for writing Gbox plug-in are on the project website: http://garmiregroup.org/granatumx/app. Additionally, we created a slack group named “GranatumX-Developer” to facilitate plug-in development from the 3rd party.

## Results

### Overview of GranatumX

The objective of GranatumX is to provide scRNA-seq biologists better access to bioinformatics tools and the ability to conduct single-cell data analysis independently (**Figure 1**). Currently other scRNA-seq platforms usually only provide a fixed set of methods implemented by the authors themselves. It is difficult to add new methods developed by the community due to programming language lock-in as well as monolithic code architectures. If a pipeline is assembled between heterogeneous tools, it is manually crafted and inhibits a repeatable execution of data analysis tools by other wet-lab scientists. As a solution, GranatumX uses the plugin and virtualized framework that provides an easy and unified approach to add new methods in a data-analysis pipeline. The plugin system is agnostic to developer code and the choice of the original scripting language. It also eliminates inter-module incompatibilities, by isolating the dependencies of each module (**Figure 2A**). As a data portal, GranatumX provides a graphical user interface (GUI) that requires no programming experience.

**Figure 1:**
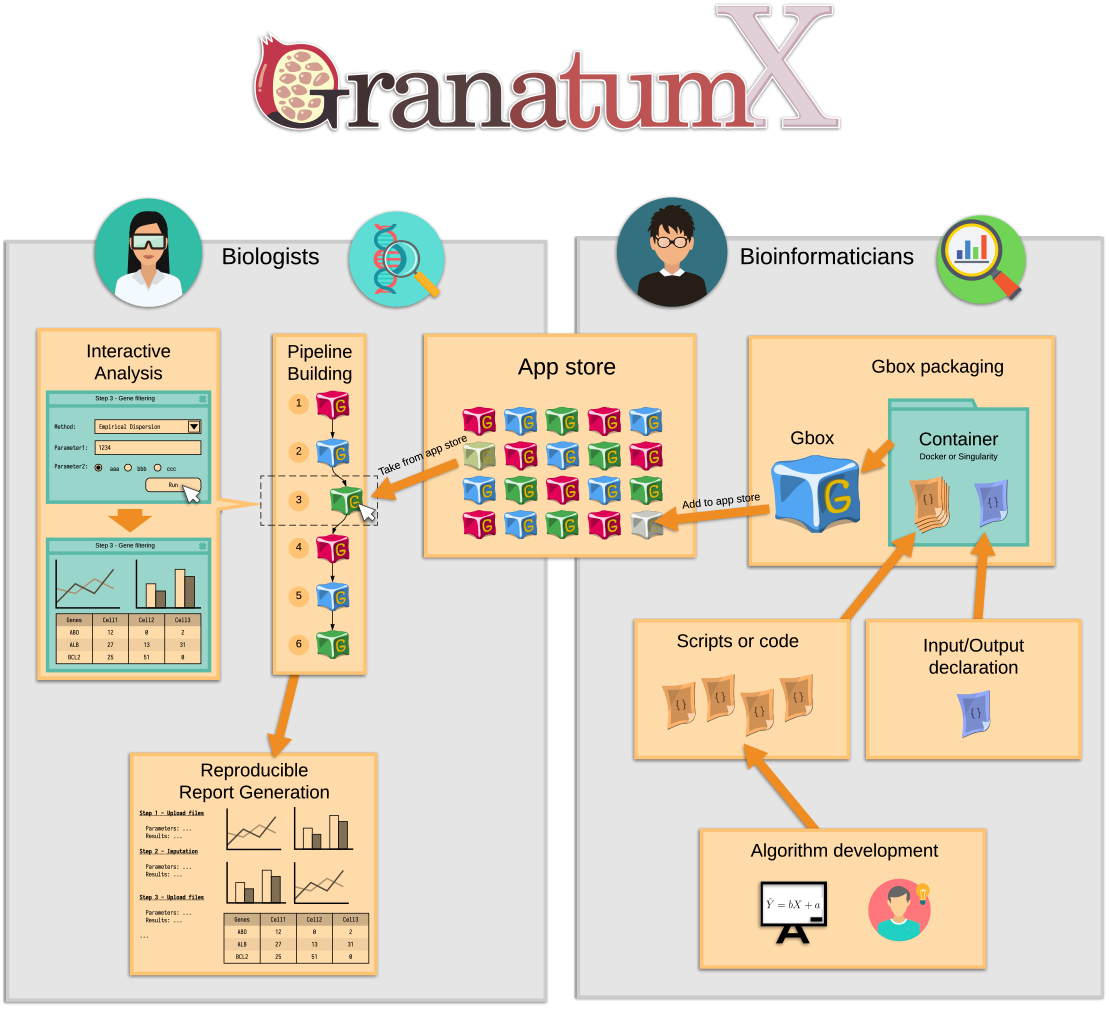
Overview of the Granatum × platform. Granatum × aims to bridge the gap between the computational method developers (the bioinformaticians) and the experiment designers (the biologists). It achieves this by building end-to-end infrastructure including the packaging and containerization of the code (**Gbox Packaging**), organization and indexing of the Gboxes (**Apps**), customization of the analysis steps (**Pipeline building**), visualization and results downloading (**Interactive Analysis**), and finally the aggregation and summarization of the study (**Report Generation**).

**Figure 2:**
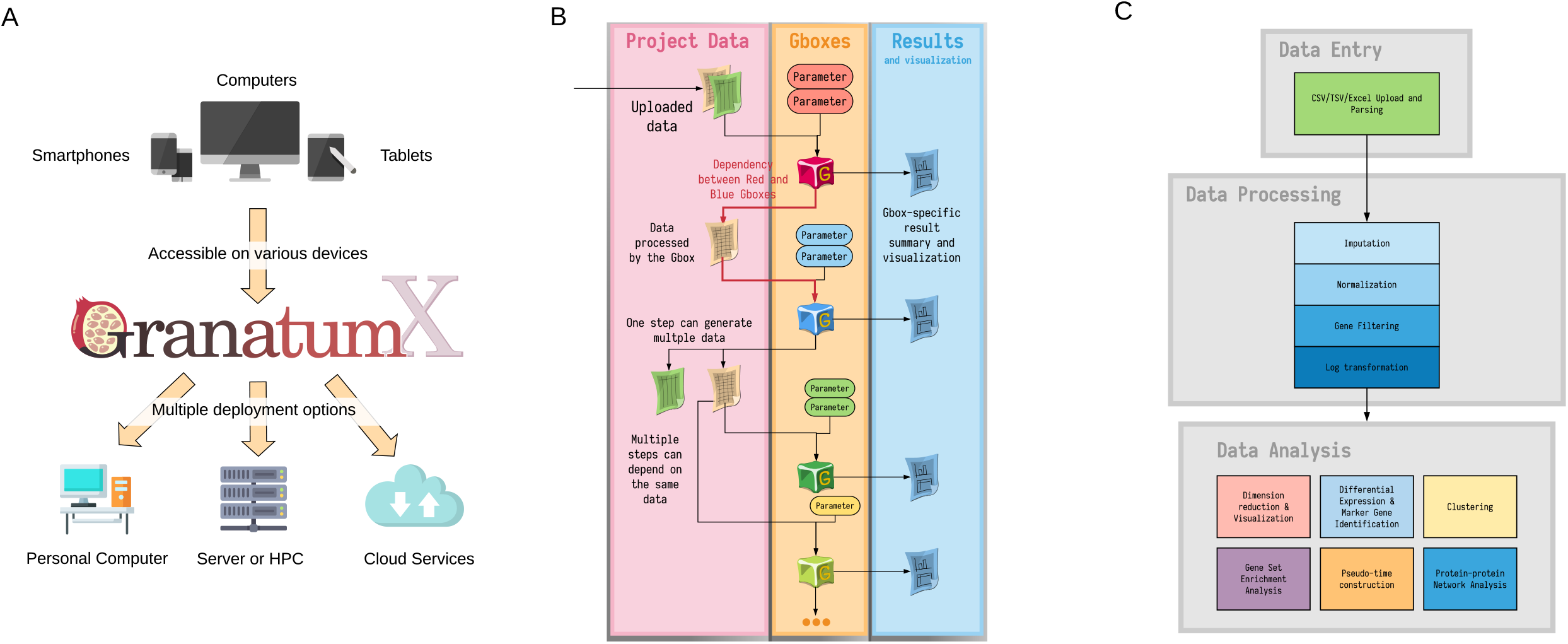
GranatumX deployment, data management and analysis flow. A) Granatum × can be deployed on various computational environments, from personal computers, private servers, High-Performance Computation systems, to cloud services. Granatum X’s web UI is adaptable to devices with various screen sizes, which allows desktop and mobile access. B) Granatum X’s data management. Each Gbox (labeled by a particular color to represent a certain functionality) with order dependency on the pipeline, may take some project data and some user-specified parameters as input and may generate results (interactive visualization, plots, tables, or even plain text) and new project data. All project data and results, as well as the specified parameters, are recorded and saved into the central data storage and can be used for reproducibility control. C) A scRNA-Seq computational study typically consists of three phases: the uploading and parsing of the expression matrices and metadata (Data Entry), the quality improvement and signal extraction of the data (Data Processing), and finally the assorted analyses on the processed data which offer biological insights (Data Analysis).

### Deployment of GranatumX

The web-based GUI can be accessed on various devices including desktop, tablets, and smartphones (**Figure 2A**). In addition to the web-based format, GranatumX is also deployable on a broad variety of computational environments, such as private PCs, cloud services, and High-Performance Computing (HPC) platforms with minimal effort by system administrators. The deployment process is unified on all platforms because all components of GranatumX are containerized in Docker [13] (also portable to Singularity [14]). GranatumX can handle larger-scale scRNA-seq datasets coming online, with an adequate cloud configuration setup and appropriate Gboxes. For example, after uploading data, it took GranatumX ~12 minutes to finish the recommended pipeline with xxx modules on an AMD 3950x with 16 cores and 128GB of DRAM memory running Ubuntu 20.04, using 10K cells downsampled from the dataset of “1.3 Million Brain Cells from E18 Mice” on the 10x Genomics website. The most time-consuming step is imputation using neural-network model DeepImpute (~⅖ time), and the detailed breakdown of time consumption is shown in **Supplementary Table 1**.

**Table 1.** Gene Set Enrichment Analysis results on clusters from UMAP plot in Figure 3B.

| Comparing<br>pairs | Gene Set Name | Gene Set Size | Normalized<br>Enrichment<br>Score | P Value | False Detection<br>Rate |
| --- | --- | --- | --- | --- | --- |
| Cluster 2 vs. Rest | KEGG<br>Glycolysis<br>Gluconeogenesis | 13 | 4.23 | 0 | 0 |
|  | KEGG<br>Pathogenic<br>Escherichia Coli<br>Infection | 14 | 3.63 | 0 | 0 |
|  | KEGG<br>Alzheimers<br>Disease | 20 | 3.97 | 0 | 0 |
|  | KEGG Tight<br>Junction | 16 | 3.03 | 0.004 | 0.0063 |
| Cluster 3 vs. Rest | KEGG Oocyte<br>Meiosis | 15 | 3.75 | 0 | 0 |
|  | KEGG<br>Pathogenic<br>Escherichia Coli<br>Infection | 14 | 3.48 | 0.001 | 0.0315 |
|  | KEGG Cell<br>Cycle | 20 | 3.13 | 0.002 | 0.042 |
|  | KEGG Ubiquitin<br>Mediated<br>Proteolysis | 12 | 3.29 | 0.005 | 0.0787 |
| Cluster 4 vs. Rest | KEGG<br>Spliceosome | 13 | 3.84 | 0 | 0 |
|  | KEGG Viral<br>Myocarditis | 25 | 3.23 | 0.002 | 0.042 |
| Cluster 5 vs. Rest | KEGG<br>Alzheimers<br>Disease | 20 | 3.08 | 0 | 0 |
|  | KEGG Antigen<br>Processing and<br>Presentation | 30 | 2.51 | 0.003 | 0.0472 |
|  | KEGG MAPK<br>Signaling<br>Pathway | 45 | 2.91 | 0 | 0 |
|  | KEGG<br>Glycolysis<br>Gluconeogenesis | 13 | 2.49 | 0.043 | 0.198 |
|  | KEGG<br>Spliceosome | 13 | 2.99 | 0.004 | 0.0504 |

### Unique Gbox modules

Gbox is a unique concept of GrantumX. It represents a containerized version of a scientific package that handles its input and output by a format understood by the GranatumX core (**Figure 2B**). GranatumX has a set of pre-installable Gboxes that enable complete scRNA-seq analysis out of the box. Various Gboxes for data entry, preprocessing, and processing can be customized and organized together, to form a complete analysis pipeline (**Figure 2C)**. One highlight feature of the Gbox is that it stands alone and the user can assume any Gbox without the need to restart the full pipeline, in case one implemented by the user fails. Another highlight of the Gbox feature is that the entire GranatumX platform is fully interactive, with addition or removal of some Gboxes or parameter changes on the go, while some other Gboxes are being executed.

A comprehensive set of over 30 Gboxes are implemented in GranatumX to perform tasks all the way from data entry and processing to downstream functional analysis. The data processing tasks help to minimize the biases in the data and increase the signal-to-noise ratio. For each of these quality improvement categories, GranatumX provides multiple popular methods from which users can pick. To assist functional analysis, GranatumX provides a core list of methods for dimension reduction, visualization (including PCA, t-SNE, and UMAP), clustering, differential expression, marker gene identification, Gene Set Enrichment Analysis, network analysis and pseudo-time construction. Versioning for each of these Gboxes has been implemented so that users can use a specific tested version of a Gbox. Developers on the other hand can work on newer versions separately before the official upgrade. Gboxes can be stored on DockerHub for public use which maintains its own versioning system (https://hub.docker.com/u/granatumx). Detailed step-by-step tutorials for writing and building Gboxes are on GranatumX website http://garmiregroup.org/granatumx/app.

### Input files

The input files of GranatumX include expression matrices and optional sample metadata tables, acceptable in a variety of formats such as CSV, TSV, or Excel format. GranatumX even accepts zip files and gz files (GNU zip), and the user can choose that format for large expression matrices. Expression matrices are raw read counts for all genes (rows) in all cells (columns). The sample metadata tables annotate each cell with a pre-assigned cell type, state, or other quality information. The parsing step creates a sparse matrix using the coordinate list (COO) format, and this representation ensures swift upload onto the back end, even for large input datasets (>10K cells). Such information will either be used to generate computational results (such as Gene Set Analysis) or be mapped onto the PCA, t-SNE, or UMAP plot for visualization (see **Figure 2C** for the workflow). Once the user uploads the gene expression matrix, the data are read into a dataframe using *Pandas* and the step updates the user with a “preview”, consisting of the first few rows and columns of the gene expression matrix, along with the number of genes and samples present.

### User-centric design

As a user-friendly tool, GranatumX allows multiple users to be affiliated with the same project for data and result sharing, while restricting one user to run the pipeline at a time to avoid data conflicts. It allows dynamically adding, removing, and reordering pipeline steps on the go. It also allows users to reset the current step. All relevant data in the analysis pipeline and all results generated by each module are stored in a database, allowing users to access and download them. To ensure reproducibility, GranatumX can automatically generate a human-readable report detailing the inputs, running arguments, and the results of all steps (see examples in **Supplementary File 1** and **Supplementary File 2**). All of these features are designed with the mindset of “consumer reports” to facilitate research in experimental labs or genomics cores.

### Case studies using GranatumX

In the following section, we demonstrate two case studies of GranatumX. The first data set was downloaded from GSE117988, including 7431 single cells generated by the 10x Genomics 3’ Chromium platform. It was obtained from a patient with metastatic Merkel cell carcinoma, treated using T cell immunotherapy as well as immune-checkpoint inhibitors (anti-PD1 and anti-CTLA4) but later developed resistance [15]. We used a customized pipeline to analyze the scRNA-seq data (**Figure 3A**). The pipeline comprises all common analysis steps, including 1) File upload, 2) imputation (based on DeepImpute [6]), 3) normalization, 4) gene filtering, 5) log transformation, 6) principal component analysis (PCA), 7) t-SNE/UMAP plot, 8) sample coloring, 9) clustering, 10) marker gene identification, 11) GSEA analysis, and 12) pseudo-time construction. The analysis report of the entire pipeline is included as **Supplementary File 1**. The clustering step identifies 7 clusters on the UMAP plot (**Figure 3B**). The exemplary GSEA analysis results (**Table 1)** show the significance in many important immune-related pathways, including the MAPK signaling pathway and antigen processing and presentation pathway (cluster_5 vs. rest), cell cycle genes (cluster_3 vs. rest), and ubiquitin-mediated proteolysis (cluster_3 vs. rest).

**Figure 3:**
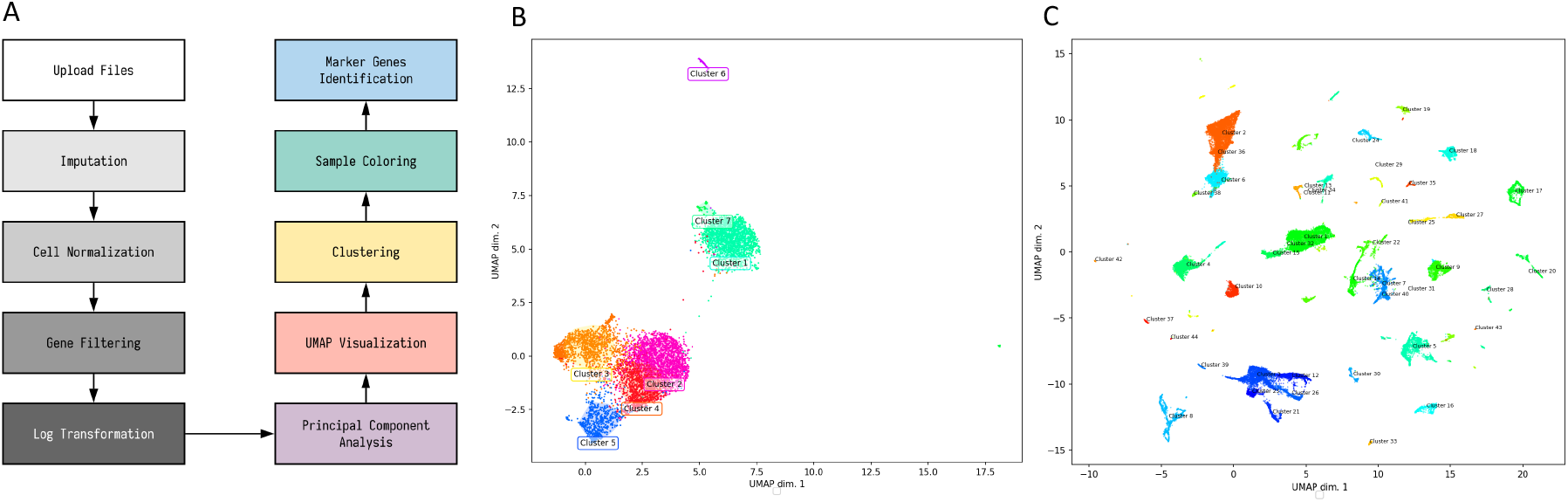
Case studies using an exemplary workflow of GranatumX. A) An exemplary workflow of a customized scRNA-seq pipeline set by the user. B) The UMAP plot clusters on Merkel cell carcinoma data from the 10x genomics platform. C) The UMAP plot clusters of Tabula Muris Consortium data.

We also used GranatumX to analyze Tabula Muris dataset, which contains 54,865 cells from 20 organs and tissues of mouse [16]. Again we used the same pipeline as shown in **Figure 3A**. GranatumX offers multiple popular clustering algorithms, and for this dataset we used the Louvain algorithm. For illustration purposes, we focus on the viewing and clustering of this large graph-based clustering method implemented by Scanpy. A total of 44 clusters are assigned on the UMAP plot (**Figure 3C**). We also superimposed the metadata that contain tissue types for each cell on the same plot for visualization (**Supplementary Figure 1**). The complete analysis report of this dataset is included as **Supplementary File 2**.

## Discussion

With the ever-increasing popularity of scRNA-seq, more and more experimental biologists will adopt this technology. At the same time, new bioinformatics tools are being developed rapidly. The development of GranatumX fills in a unique niche that enables both scientific and technical advancements. It is a “common ground” that connects scRNA-seq tool developers with the end-users, together for new discoveries. Domain experts can use GranatumX for the initial exploratory analysis. Additionally, with more Gboxes to be implemented on model performance metrics, GranatumX could be a vessel to enable benchmark studies to compare existing computational modules and pipelines, as well as assess the performance of a new method or pipeline relative to the existing ones. Moreover, it can also serve as the test engine to probe the source of variations in different modules, so as to optimize a pipeline for given datasets.

To demonstrate the uniqueness of GranatumX, we also compare it with other similar tools for comprehensive scRNA-seq analysis, such as SC1[17], ASAP [10] and Single Cell Explorer [18] in **Table 2**. While all these tools aim for simple report and interaction with biologists, GranatumX is the only framework that supports bioinformatics developers to contribute their own plug-ins (**Table 2**). This significantly enhances the adaptability of GranatumX among the developer community. The web-tool that is closest to GranatumX is Single Cell Explorer, still with significant differences in the functionalities. Single Cell Explorer begins from raw data processing including reading mapping alignment. GranatumX is a much lighter-weight tool that starts from a cell read count table since the alignment/tag counting step is readily done by the popular Cell Ranger software of the 10x genomics platform. Instead, GranatumX put more efforts on downstream analysis, such as gene enrichment analysis, protein-protein interaction and pseudo-time construction (**Table 2**). For ASAP, besides lacking modules to perform functions such as imputation, protein-protein interaction and pseudo-time construction, it also does not allow reconfiguration of the pipeline like GranatumX. SC1 lacks the flexibility and functionalities similar to ASAP and is restricted by Shiny, an R programming language based web-interface, whereas GranatumX accepted containerized Gboxes packaged written in R, Python, or other languages.

**Table 2.**
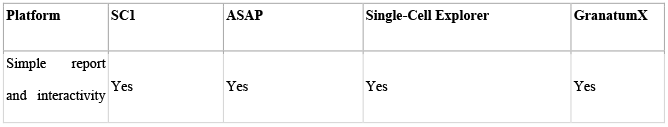
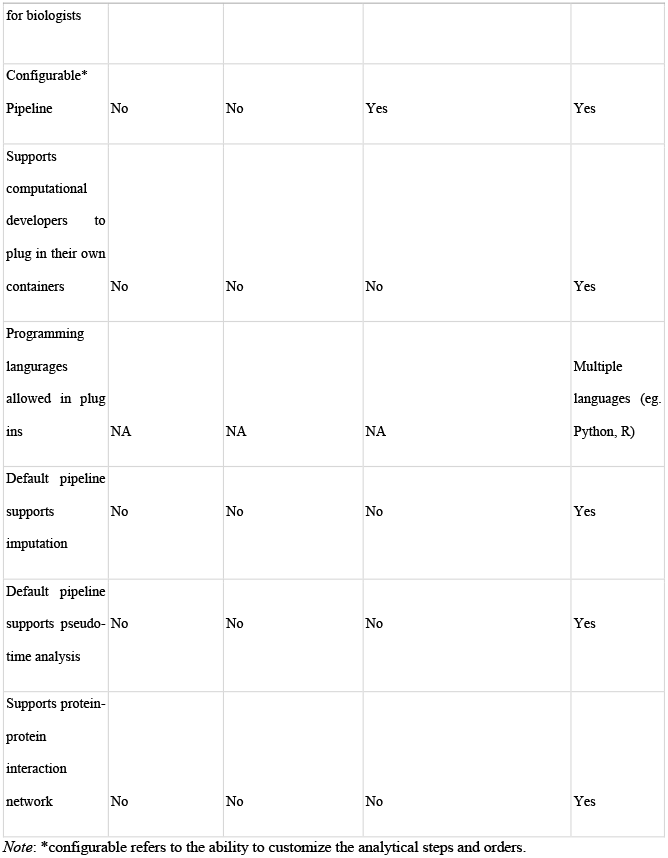
Comparison on multiple user-friendly web-tools.

As an inclusive and open software environment that employs other third-party tools, GranatumX has some challenges. One of them is handling the upgrade of underlying 3rd party libraries and resources. Accompanying with updated 3rd party tools which may not be tested extensively by the original developers, errors from these packages may propagate into GranatumX. To deal with this issue, we implement versioning through the use of Docker which helps to maintain system-level dependencies as well as software dependencies in a complete package. The Gbox Docker containers for this release are listed in the **Supplementary Table 2** with a version number of 1.0.0. New versions can update the minor and major revision numbers so that users know exactly which code is being executed for a given pipeline. The source code for the Docker containers which represent the Gboxes are stored in the corresponding GitHub repositories. For example, https://hub.docker.com/r/granatumx/gbox-differentialexpression is stored in https://github.com/granatumx/gbox-differentialexpression. Such an endeavor provides safety and reliability in maintaining the stability of the software not just in the source of the software but in the configuration of the system required to run the computational elements. Due to its openness, GranatumX can not prevent p-hacking or manipulating data analysis to improve the statistical significance of the desired result [19]. One way to discourage p-hacking is to suggest using standard pipeline and default parameters. If the user chooses values other than defaults, the reproducible design of GranatumX allows one to compare the outputs from the users (if they are recorded) with those from the default setting.

## Conclusions

We present an open-source, shareable and evolvable single cell analysis tool called GranantumX. It not only enables the domain experts to independently conduct single cell analysis, but also promotes bioinformatics tool developers to contribute and develop their own single cell analysis methods through Gbox plug-in setup. We hope that GranatumX will engage the single cell analysis community broadly and continuously for scientific discoveries.

## Code availability

The webtool of GranatumX can be found at http://garmiregroup.org/granatumx/app. On this website, users can also find youtube or youku tutorial videos that demonstrate how to use GranatumX webtool. The source code for GranatumX is available at https://github.com/granatumx under MIT license. All builds are deployed via Docker Hub at https://hub.docker.com/u/granatumx.

## Copyright

Some of the cartoon icons in Figure 1 and Figure 2 are downloaded from https://www.flaticon.com/.

## CRediT authorship contribution statement

Lana × Garmire: Conceptualization, supervision, writing manuscript. David Garmire, Xun Zhu: Design, implementation of Software, writing manuscript. Aravind Mantravadi, Breck Yunits: Implementation and code testing. Qianhui Huang, Yu Liu: case study and testing. Thomas Wolfgruber, Olivier Poirion, Tianying Zhao, Cédric Arisdakessian, Stefan Stanojevic: implementation of Gbox. All authors have read and approved the manuscript.

## Acknowledgment

We thank Dr. Larry Reiter for providing valuable feedback while using GrantumX to test his single cell dataset. This research was supported by grants K01ES025434 awarded by NIEHS through funds provided by the trans-NIH Big Data to Knowledge (BD2K) initiative (www.bd2k.nih.gov), P20 COBRE GM103457 awarded by NIH/NIGMS, R01 LM012373 awarded by NLM, R01 HD084633 awarded by NICHD to L.X. Garmire.

---

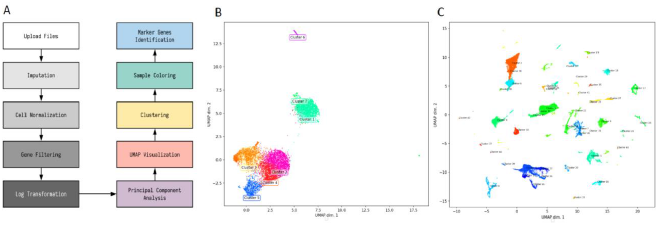

## Supplementary Materials

**Supplementary Figure 1. UMAP plot with annotated tissue types in the Tabula Muris Consortium data**.

**Supplementary Table 1. The running time for different steps of the recommended pipeline in GranatumX over 10**,**000 cells, using an AMD 3950X processor, 16 cores, with 128 GB of RAM**.

**Supplementary Table 2. The list of currently implemented Gboxes**.

**Supplementary File 1: The analysis report using a dataset with metastatic Merkel cell carcinoma from the 10x genomics platform**.

**Supplementary File 2: The analysis report using a dataset with Tabula Muris Consortium data**.

