## Supplementary figures and images for "GranatumX: A Community-engaging, Modularized, and Flexible Webtool for Single-cell Data Analysis"

### Supplementary Figure 1

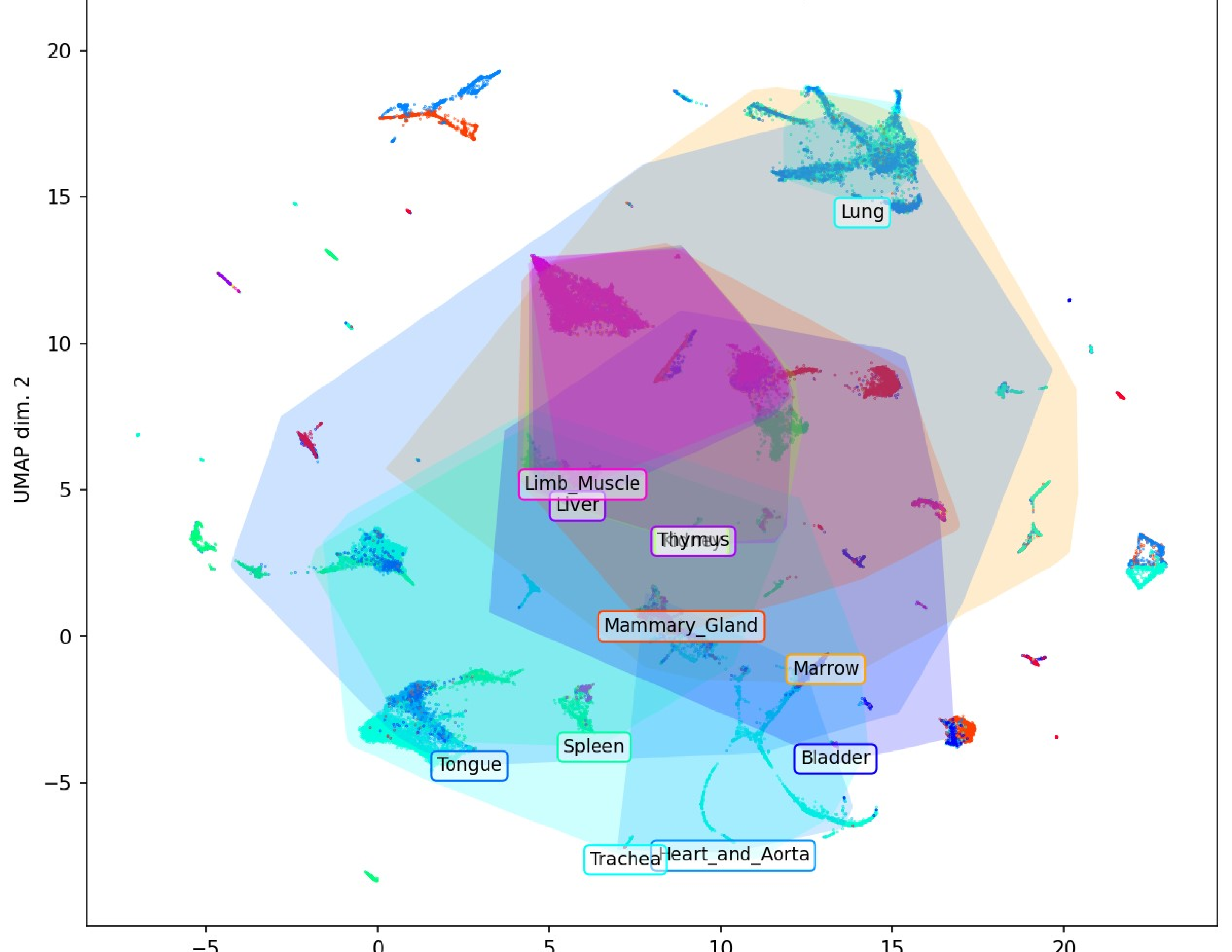
