## Supplementary material for "GranatumX: A Community-engaging, Modularized, and Flexible Webtool for Single-cell Data Analysis": Supplemetnary File 2

### Supplementary File 1

#### Analysis Report

This report is generated by GranatumX

Please cite: Zhu, Xun, et al. "GranatumX: A community engaging and flexible software environment for single-cell analysis." bioRxiv (2018): 385591.

This is the pipeline to replicate Supplementary file 1 with GSE117988 data

### Upload Files 1.0.0

Assay to upload: **GSE117988_raw.expMatrix_Tumor.csv.zip (12.91 MB)**
File format: **"zip"**
Convert gene IDs: **false**
Species: **"human"**
Convert gene IDs into (HGNC symbol is recommended): **"symbol"**
Add extra info (from BioMart) into gene metadata: **true**
Enter your email address to get notified of any errors encountered in the pipeline: **false**

The assay has **21861** genes (with inferred ID type: Symbol) and **7431** samples.

The first few rows and columns:

```
0.0, 0.0, 0.0, 0.0, 0.0, 0.0, 0.0, 0.0, 0.0, 0.0
1.0, 0.0, 0.0, 0.0, 0.0, 0.0, 0.0, 0.0, 0.0, 0.0
0.0, 0.0, 0.0, 0.0, 0.0, 0.0, 0.0, 0.0, 0.0, 0.0
0.0, 0.0, 0.0, 0.0, 0.0, 0.0, 0.0, 0.0, 0.0, 0.0
0.0, 0.0, 0.0, 0.0, 0.0, 0.0, 0.0, 0.0, 0.0, 0.0
0.0, 0.0, 0.0, 0.0, 0.0, 0.0, 0.0, 0.0, 0.0, 0.0
0.0, 0.0, 1.0, 0.0, 1.0, 0.0, 0.0, 0.0, 0.0, 0.0
0.0, 0.0, 0.0, 0.0, 0.0, 0.0, 0.0, 1.0, 0.0, 0.0
0.0, 0.0, 0.0, 0.0, 0.0, 0.0, 0.0, 0.0, 0.0, 0.0
0.0, 0.0, 0.0, 0.0, 0.0, 0.0, 0.0, 0.0, 0.0, 0.0
```

- Finished upload step in 239.38 seconds*

### DeepImpute 2.0.0

Random seed: **12345**
Use automatic gene imputation limit: **true**
Gene rank limit: **2000**
Cell subset: **1**
Assay: **[A]GSE117988_raw.expMatrix_Tumor.csv.zip** (from step 1: Upload Files 1.0.0)

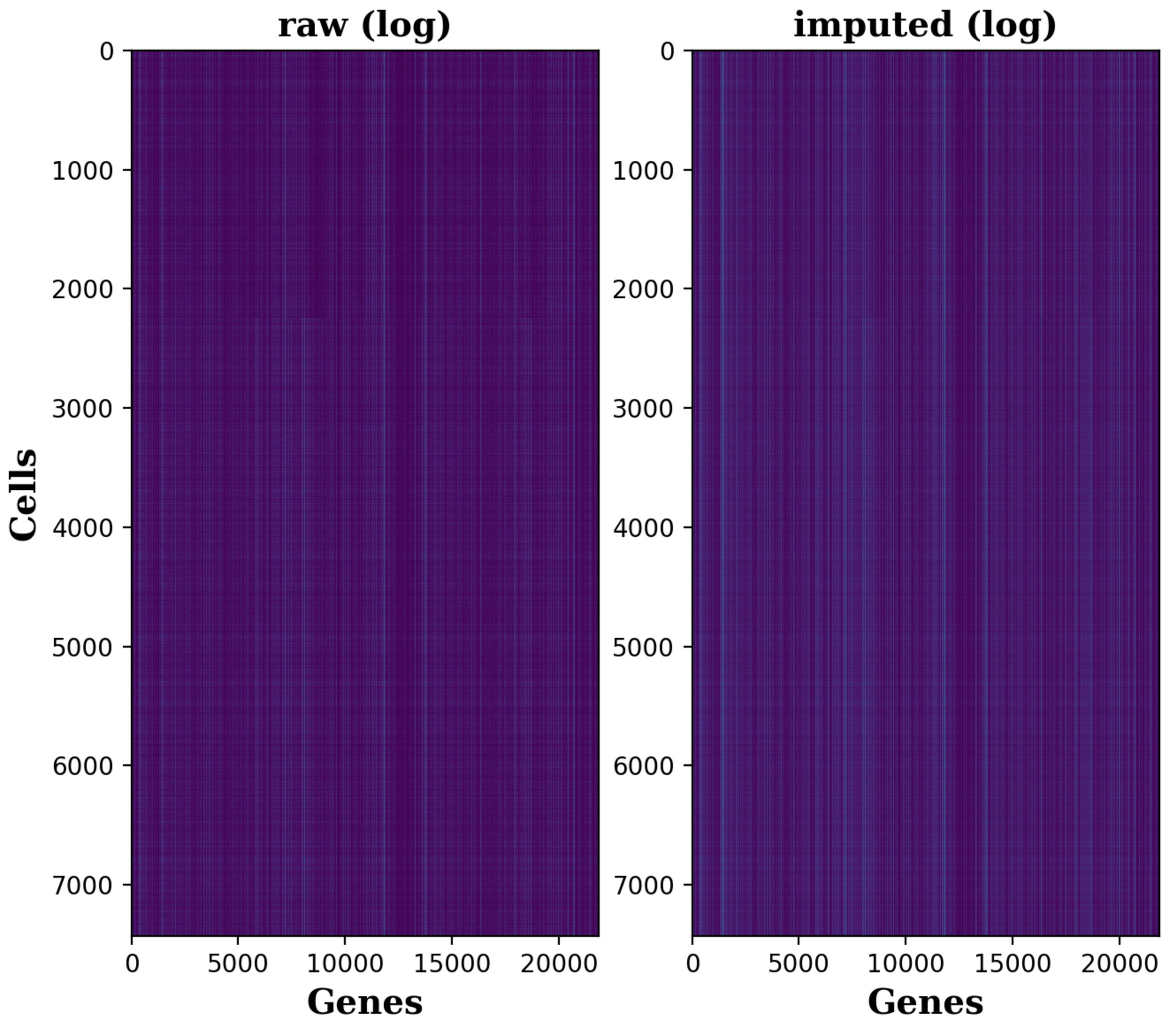


Heatmaps

- Data frame number of rows: **7431**
- Data frame number of columns: **21861**
- Number of imputed genes: **1536**

- Percentage of dropout entries *before* imputation: **92.70%**
- Percentage of dropout entries *after* imputation: **89.11%**
- Accuracy (correlation) on masked data: **0.92**

### Cell Normalization 1.0.0

Log transform in the boxplots: **false**
Normalization method: **"quantile"**
Number of cells to plot in the bar-plot: **40**
Assay: **Imputed assay** (from step 2: DeepImpute 2.0.0)

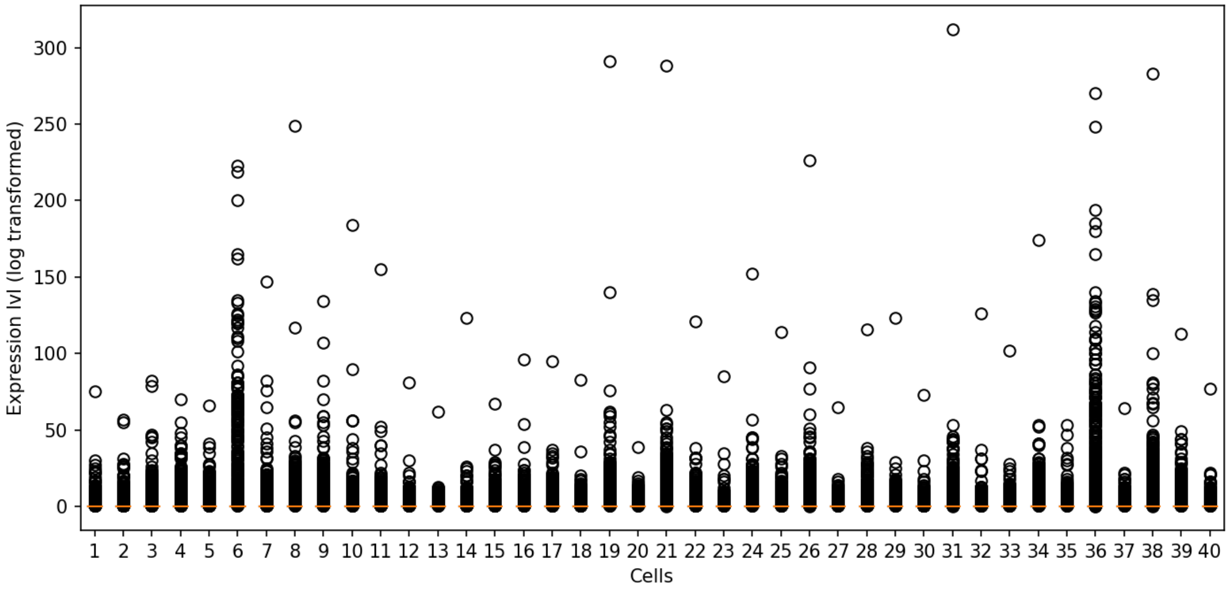


Before normalization: Each bar in the box plot represents one cell.

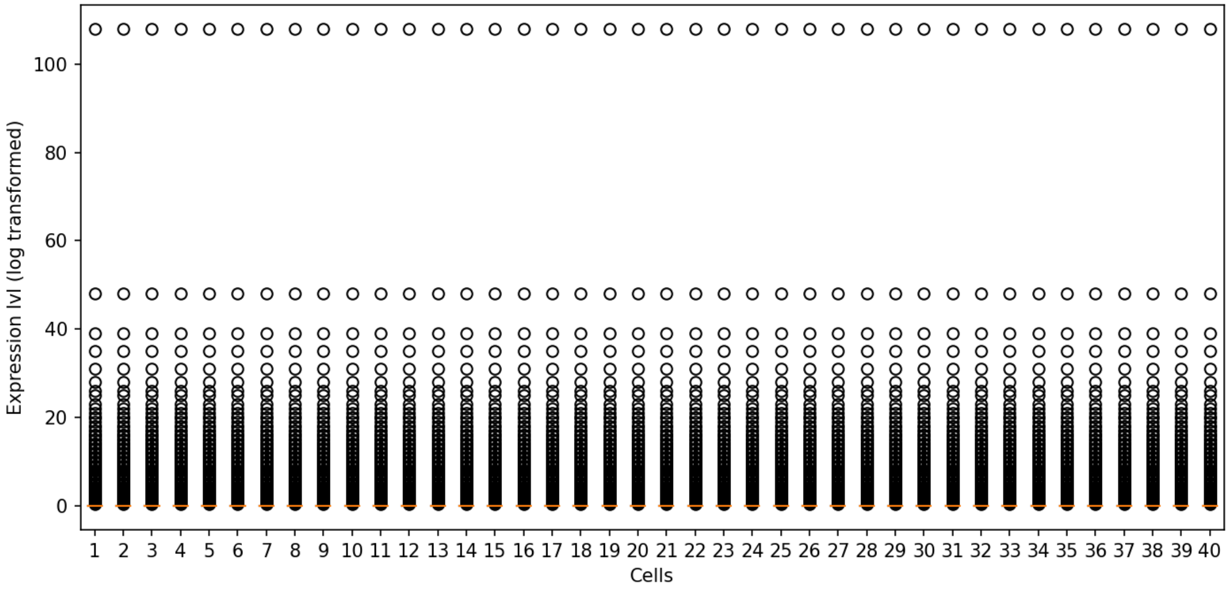


After normalization: Each bar in the box plot represents one cell.

### Scanpy Gene Filtering 1.0.0

The gene has to be expressed in at least ___ cells: **3**
The average expression level of the gene has to be greater than: **1**
The average expression level of the gene has to be less than: **999999**
The dispersion of the gene has to be greater than: **0.5**
The dispersion of the gene has to be less than: **999999**
Assay: **Normalized assay** (from step 3: Cell Normalization 1.0.0)

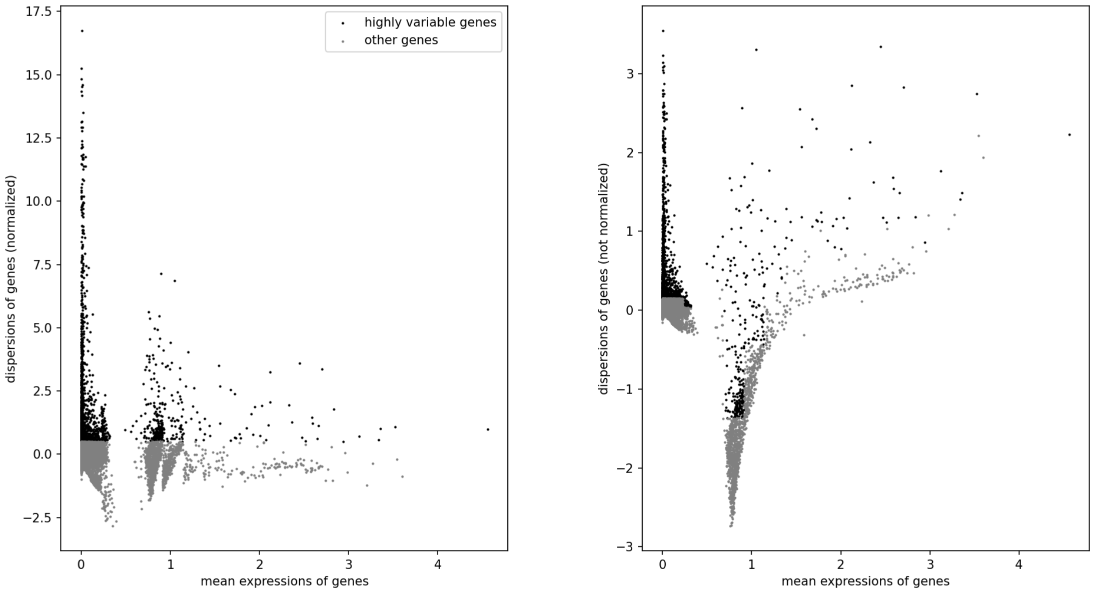


Each dot represent a gene. The gray dots are the removed genes. The x-axis is log-transformed.

Number of genes before filtering: **21861**

Number of genes after filtering: **1594**

### Log transformation 1.0.0

The base used for the log function: **2**

The pseudo counts added before log transformation (to avoid getting log(0)): **1**

Assay including matrix and geneIds: **Filtered Assay** (from step 4: Scanpy Gene Filtering 1.0.0)

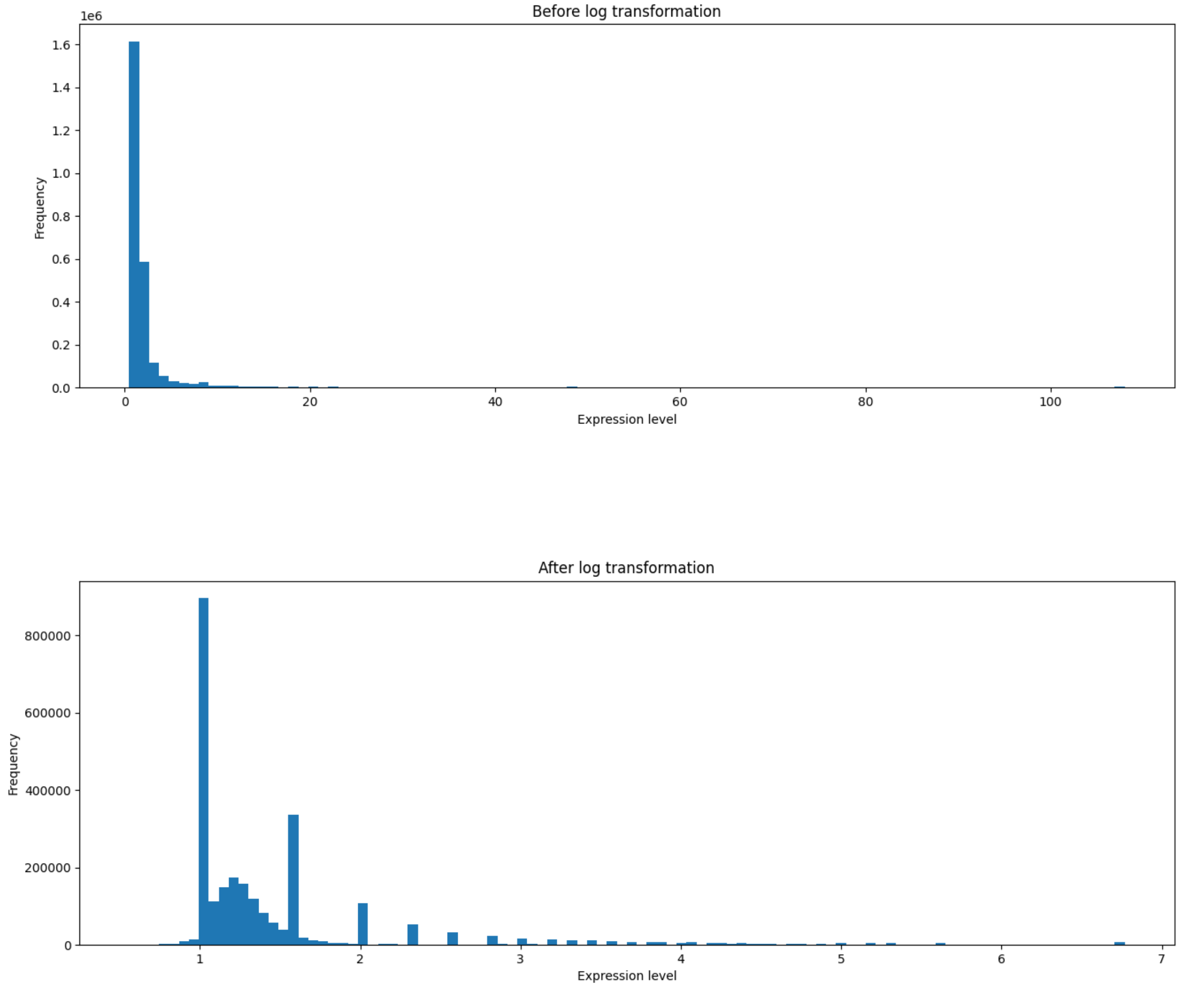


The distribution of expression level before and after log transformation. Only the values greater than the 5 percentile (usually zero in single-cell data) and lower than 95 percentile are considered.

### Principal Component Analysis 1.0.0

Number of top components to calculate: **2**

Assay: **Log transformed assay** (from step 5: Log transformation 1.0.0)

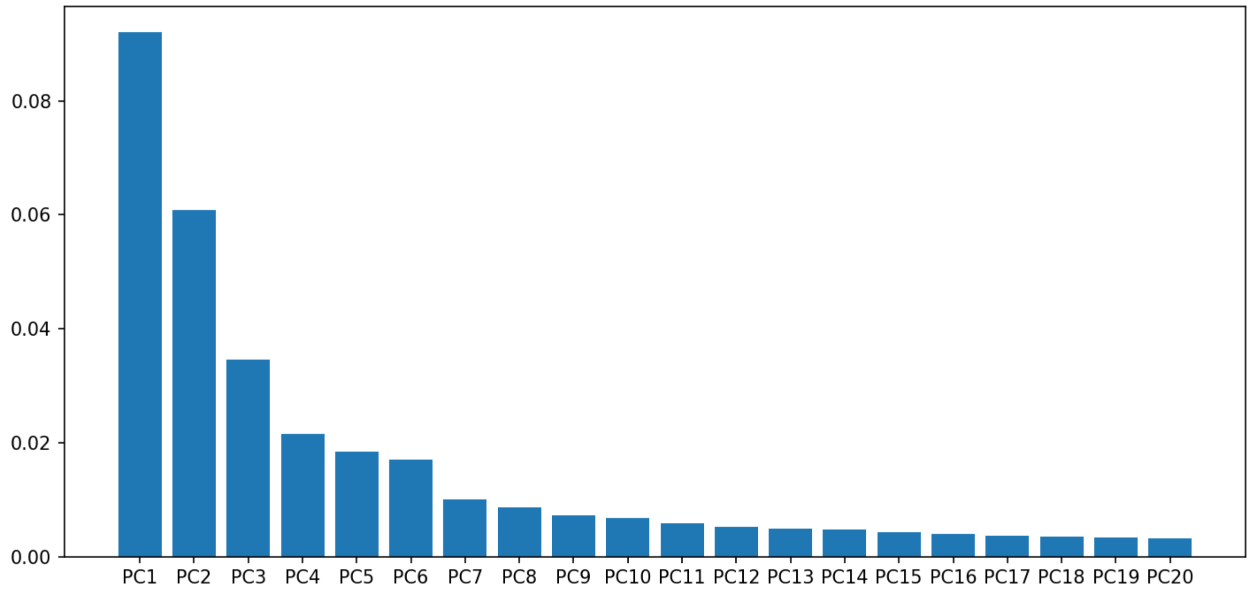


Explained variance (ratio) by each Principal Component (PC)

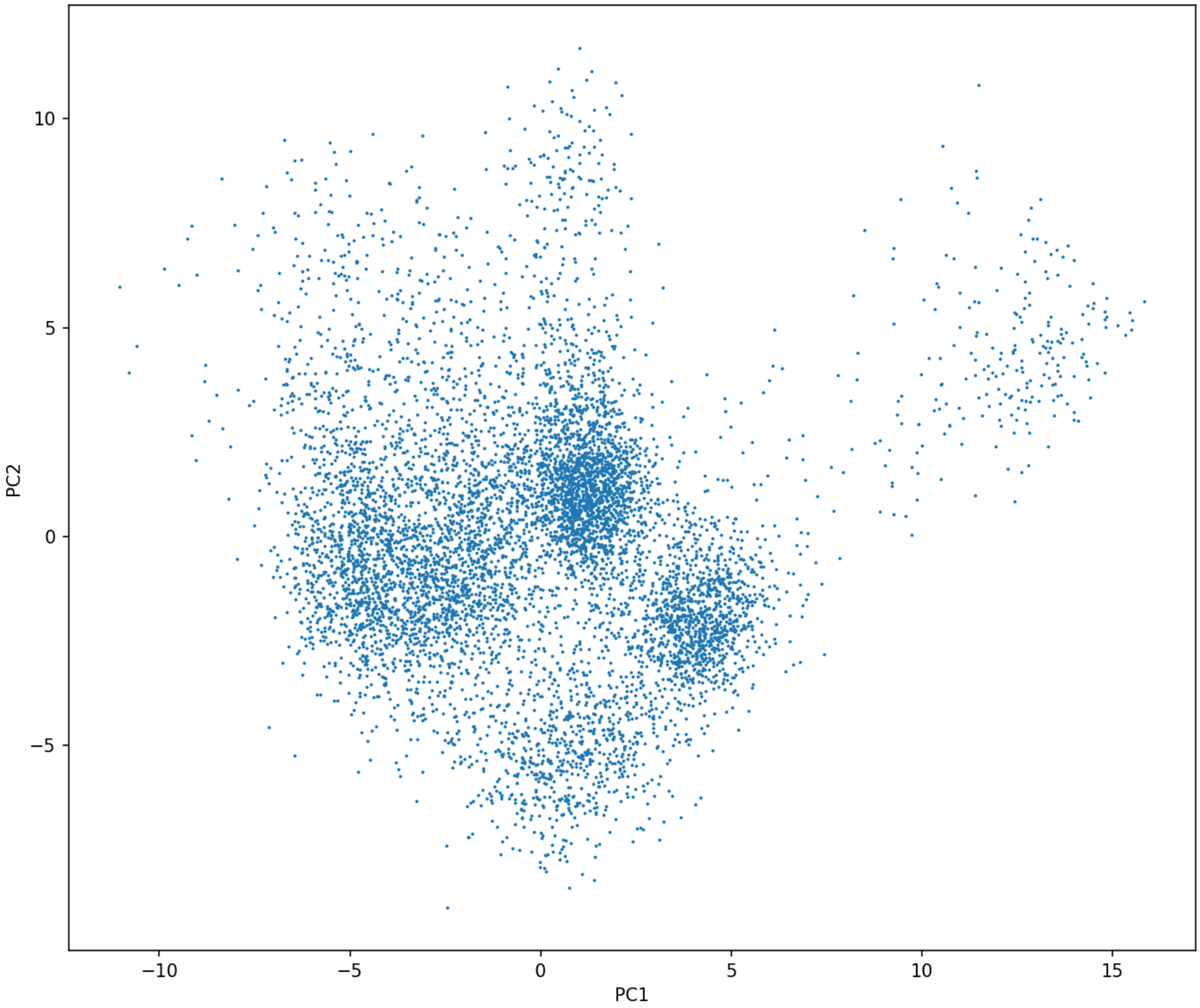


PC1 vs. PC2

### t-Distributed Stochastic Neighbor Embedding 1.0.0

Random seed: **56143**

Assay: **Log transformed assay** (from step 5: Log transformation 1.0.0)

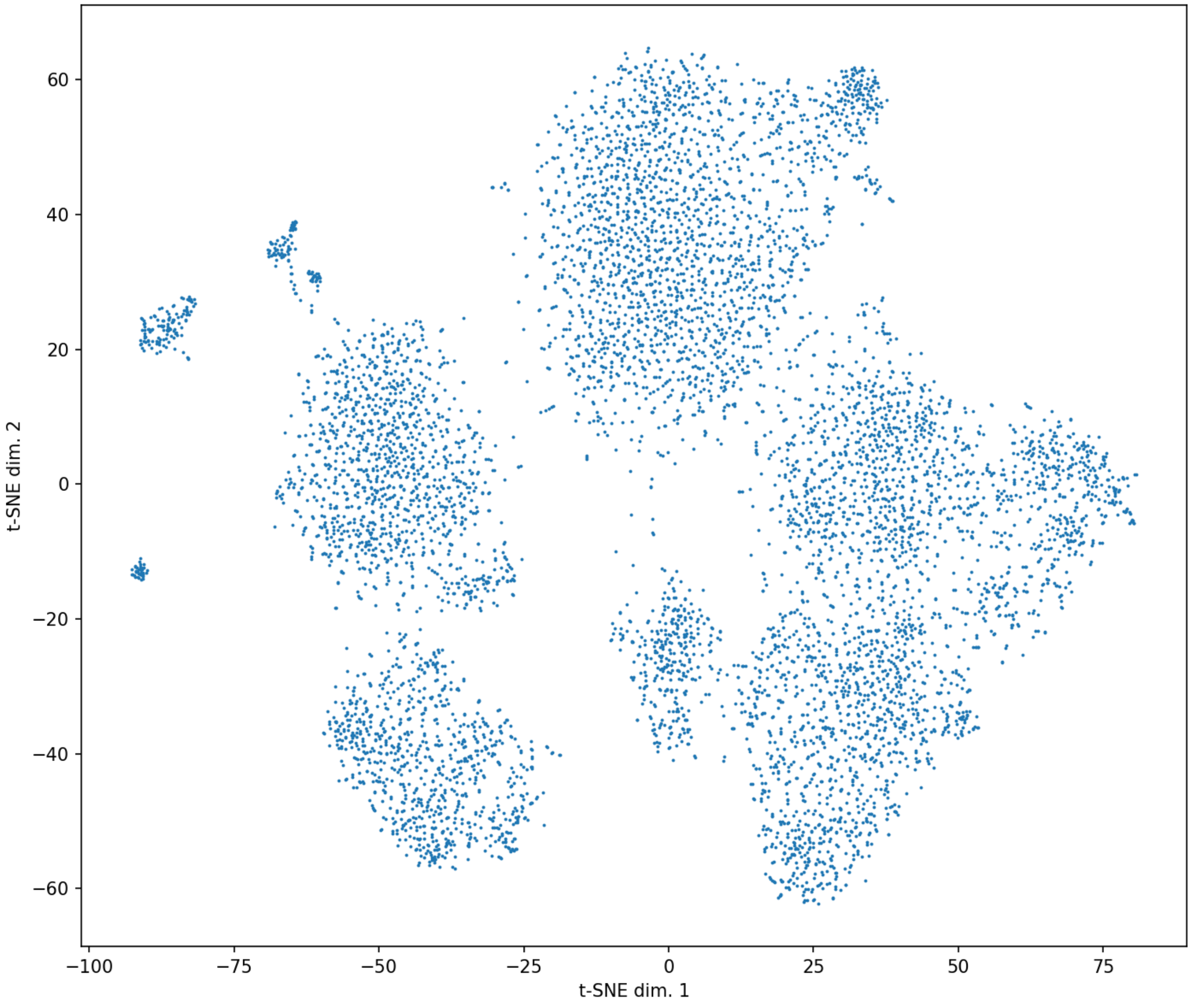


t-SNE plot: each dot represents a cell

### UMAP 1.0.0

Number of neighbors (n_neighbors, an integer): **15**
Minimum distance (min_dist, a real number ranges from 0 to 1): **0.1**
Metric (metric): **"euclidean"**
Assay: **Log transformed assay** (from step 5: Log transformation 1.0.0)

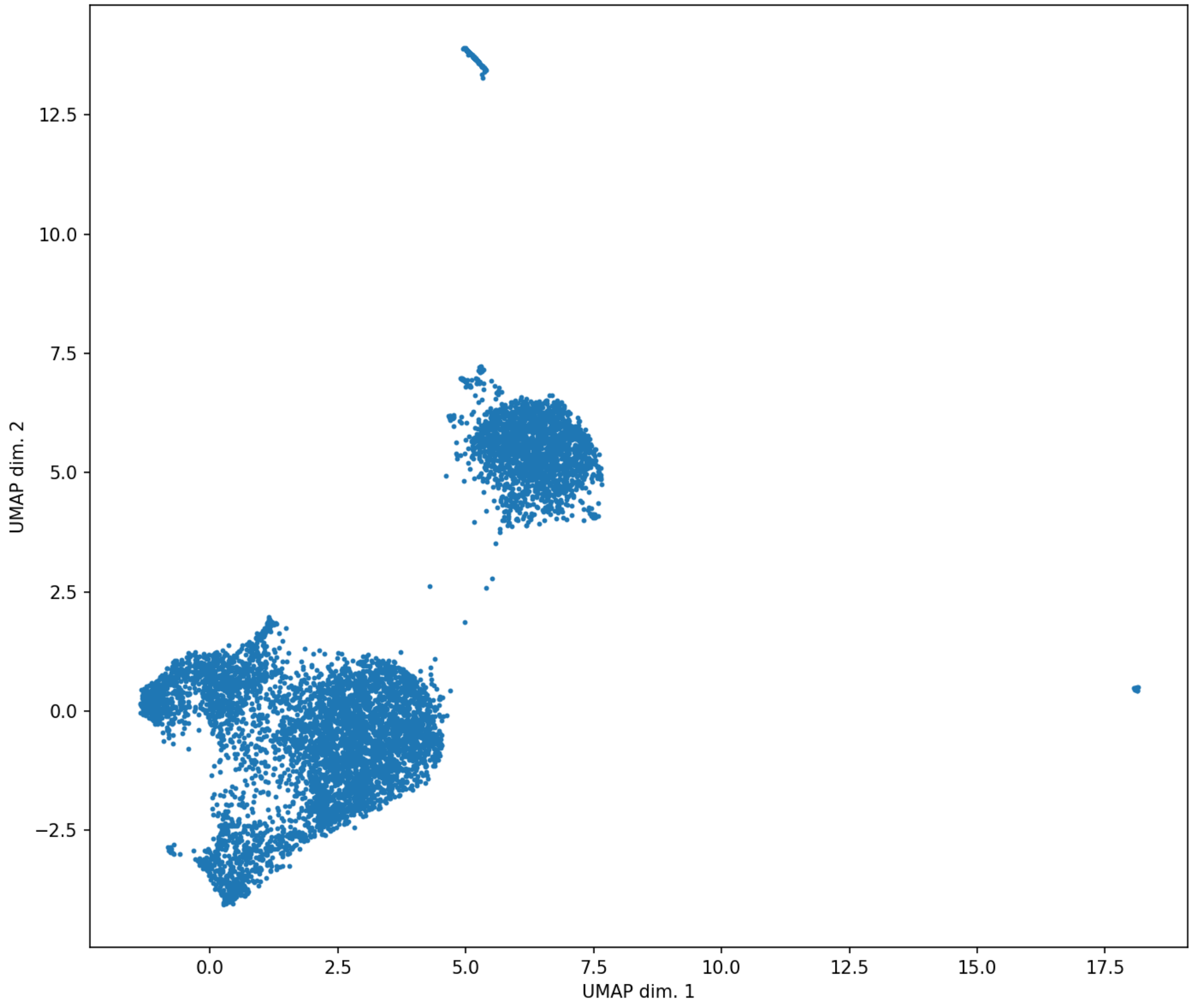


UMAP plot: each dot represents a cell

### ScanpyClustering 1.0.0

Random seed: **13513**

Assay including matrix and geneIds: **Log transformed assay** (from step 5: Log transformation 1.0.0)

Cell coordinates for visualization: **PC1 vs. PC2** (from step 6: Principal Component Analysis 1.0.0)

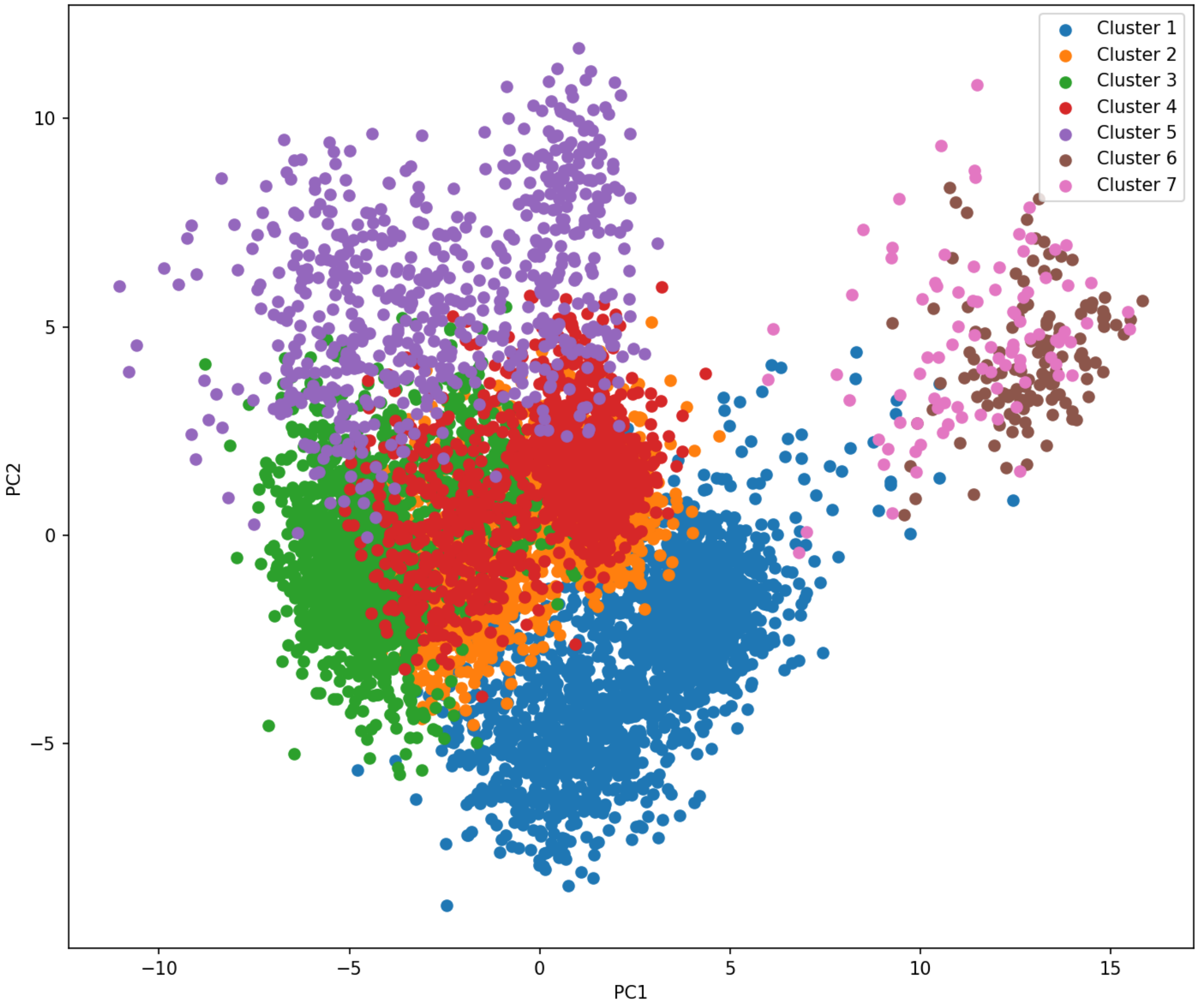


Scatter-plot using imported cell coordinates. Each dot represents a cell. The colors indicate the indentified cell clusters.

### ScanpyClustering 1.0.0

Random seed: **13513**

Assay including matrix and geneIds: **Log transformed assay** (from step 5: Log transformation 1.0.0)

Cell coordinates for visualization: **t-SNE coordinates** (from step 7: t-Distributed Stochastic Neighbor Embedding 1.0.0)

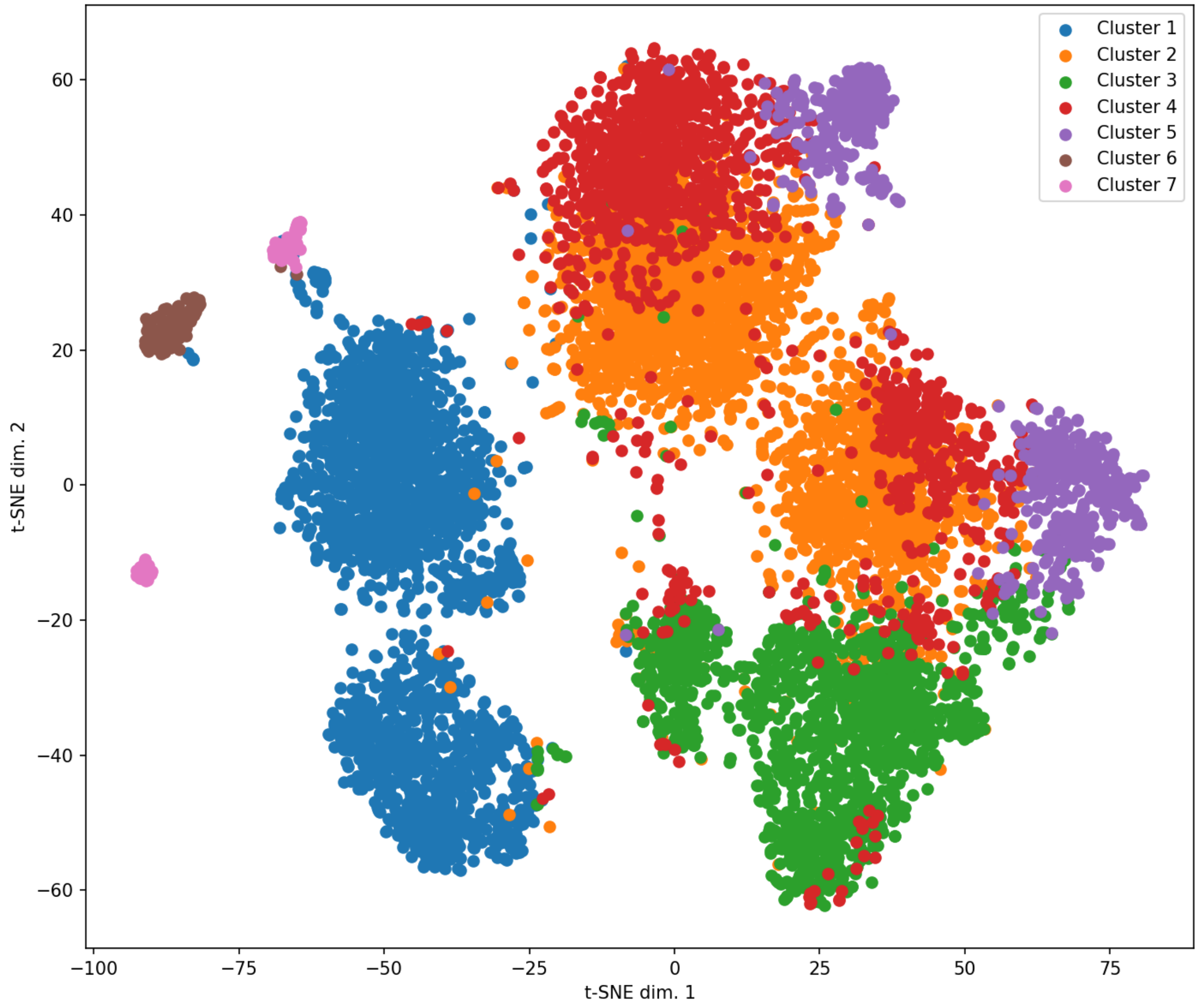


Scatter-plot using imported cell coordinates. Each dot represents a cell. The colors indicate the indentified cell clusters.

### ScanpyClustering 1.0.0

Random seed: **13513**

Cell coordinates for visualization: **UMAP coordinates** (from step 8: UMAP 1.0.0)

Assay including matrix and geneIds: **Log transformed assay** (from step 5: Log transformation 1.0.0)

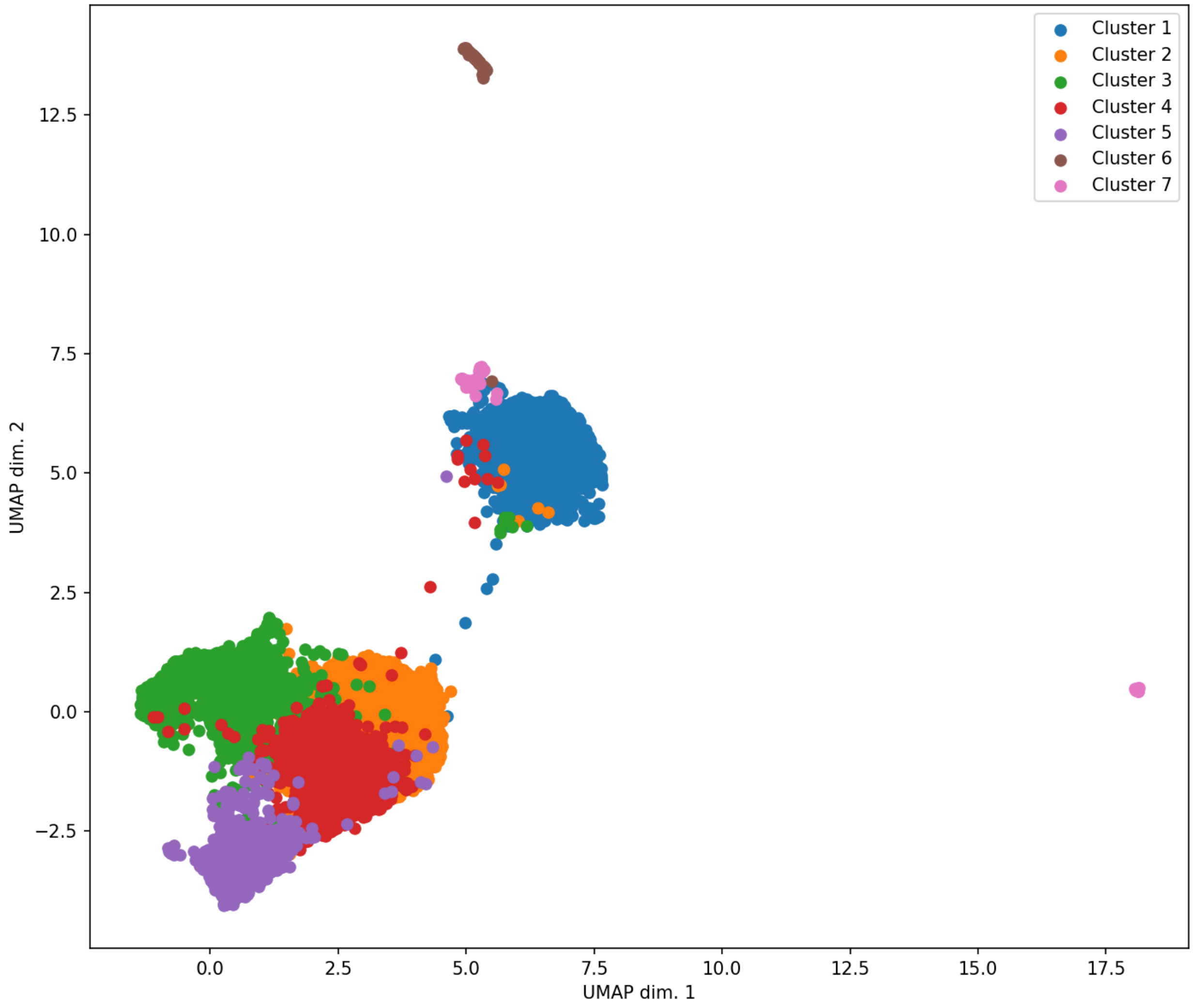


Scatter-plot using imported cell coordinates. Each dot represents a cell. The colors indicate the indentified cell clusters.

### Marker Genes Identification 1.0.0

Group vector: **Cluster assignment** (from step 11: ScanpyClustering 1.0.0)

Assay including matrix and geneIds: **Log transformed assay** (from step 5: Log transformation 1.0.0)

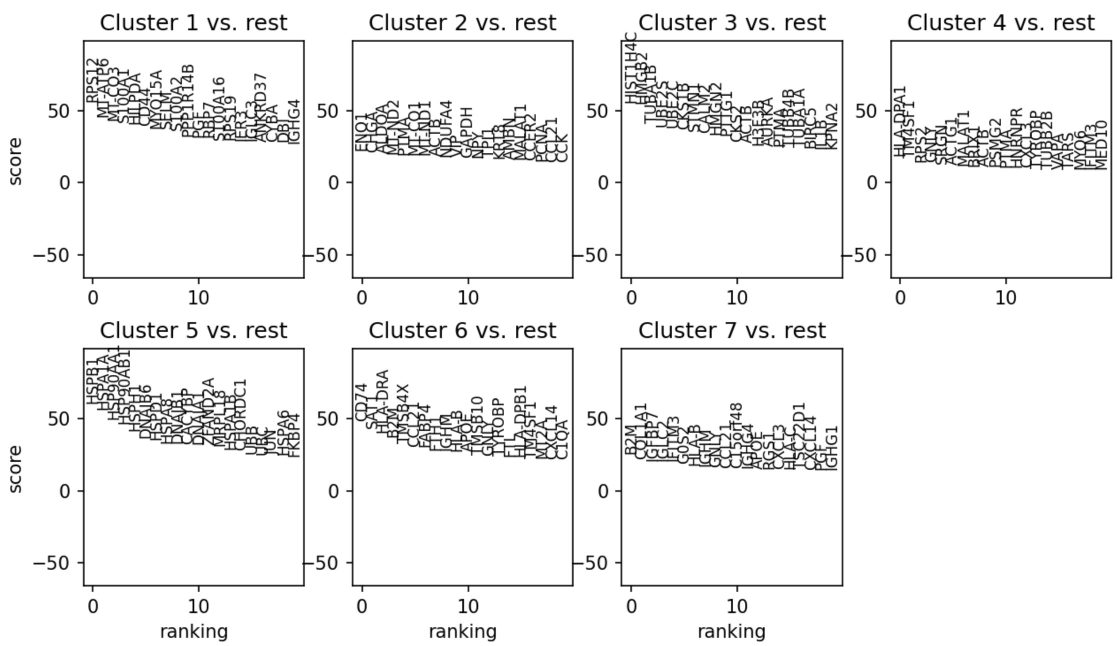


One-vs-rest marker genes

### Broad GSEA 1.0.0

The species: **"human"**
The database for the enrichement analysis: **"kegg"**
Number of repeats for calculating p-values: **1000**
A list of genes with their scores (usually output from scanpy scoring for example): **Marker score (Cluster 1 vs. rest)** (from step 12: Marker Genes Identification 1.0.0)

| gset_name | gset_size | nes | p_val | fdr |
|---|---|---|---|---|
| KEGG_ANTIGEN_PRC | 30 | 2.4183472526 | 0.001 | 0.063 |
| KEGG_MAPK_SIGNAI | 45 | 2.242569416 | 0.005 | 0.105 |
| KEGG_VIRAL_MYOCA | 25 | 2.459714285 | 0.005 | 0.105 |
| KEGG_PATHOGENIC_ | 14 | 2.5451885223 | 0.007 | 0.11025 |
| KEGG_REGULATION_ | 25 | 2.3859803876 | 0.01 | 0.126 |
| KEGG_GLYCOLYSIS_G | 13 | 2.3783434423 | 0.021 | 0.189 |
| KEGG_TYPE_I_DIABE | 17 | 2.3151470657 | 0.021 | 0.189 |
| KEGG_TIGHT_JUNCT | 16 | 2.3476164319 | 0.035 | 0.275625 |
| KEGG_GRAFT_VERSU | 19 | 2.1367696417 | 0.047 | 0.2953125 |

### Broad GSEA 1.0.0

The species: **"human"**
The database for the enrichement analysis: **"kegg"**
Number of repeats for calculating p-values: **1000**
A list of genes with their scores (usually output from scanpy scoring for example): **Marker score (Cluster 2 vs. rest)** (from step 12: Marker Genes Identification 1.0.0)

| gset_name | gset_size | nes | p_val | fdr |
|---|---|---|---|---|
| KEGG_GLYCOLYSIS_G | 13 | 4.2332424448 | 0 | 0 |
| KEGG_PATHOGENIC_ | 14 | 3.6287072467 | 0 | 0 |
| KEGG_ALZHEIMERS_ | 20 | 3.969936 | 0 | 0 |
| KEGG_TIGHT_JUNCT | 16 | 3.0309404996 | 0.004 | 0.063 |
| KEGG_HUNTINGTON: | 16 | 2.8748446055 | 0.009 | 0.1134 |
| KEGG_REGULATION_ | 25 | 2.3360851389 | 0.036 | 0.324 |
| KEGG_PARKINSONS_ | 12 | 2.5324904944 | 0.036 | 0.324 |
| KEGG_UBIQUITIN_ME | 12 | 2.3614030539 | 0.054 | 0.42525 |
| KEGG_OOCYTE_MEIC | 15 | 2.1520125214 | 0.08 | 0.56 |

### Broad GSEA 1.0.0

The species: **"human"**
The database for the enrichement analysis: **"kegg"**
Number of repeats for calculating p-values: **1000**
A list of genes with their scores (usually output from scanpy scoring for example): **Marker score (Cluster 3 vs. rest)** (from step 12: Marker Genes Identification 1.0.0)

| gset_name | gset_size | nes | p_val | fdr |
|---|---|---|---|---|
| KEGG_OOCYTE_MEIC | 15 | 3.9142438048 | 0 | 0 |
| KEGG_PATHOGENIC_ | 14 | 3.6276032782 | 0.001 | 0.021 |
| KEGG_UBIQUITIN_ME | 12 | 3.2339121482 | 0.001 | 0.021 |
| KEGG_CELL_CYCLE | 20 | 3.0585714514 | 0.004 | 0.063 |
| KEGG_REGULATION_ | 25 | 2.3460916724 | 0.039 | 0.35 |
| KEGG_ANTIGEN_PRC | 30 | 2.2842579982 | 0.04 | 0.35 |
| KEGG_INSULIN_SIGN | 12 | 2.5842161078 | 0.05 | 0.35 |
| KEGG_TIGHT_JUNCT | 16 | 2.4147621326 | 0.05 | 0.35 |
| KEGG_ALLOGRAFT_R | 17 | 2.3470481376 | 0.06 | 0.35 |

### Broad GSEA 1.0.0

The species: **"human"**
The database for the enrichement analysis: **"go"**
Number of repeats for calculating p-values: **1000**
A list of genes with their scores (usually output from scanpy scoring for example): **Marker score (Cluster 1 vs. rest)** (from step 12: Marker Genes Identification 1.0.0)

| gset_name | gset_size | nes | p_val | fdr |
|---|---|---|---|---|
| GO_CONDENSED_CH | 17 | 2.7298822377 | 0 | 0 |
| GO_REGULATION_OF | 139 | 1.9514943306 | 0 | 0 |
| GO_RNA_BINDING | 138 | 2.0809844395 | 0 | 0 |
| GO_CONDENSED_CH | 24 | 2.8174020056 | 0 | 0 |
| GO_ESTABLISHMENT | 152 | 1.8743619136 | 0 | 0 |
| GO_REGULATION_OF | 23 | 2.7517004946 | 0 | 0 |
| GO_MITOTIC_SISTER | 16 | 2.6126316537 | 0 | 0 |
| GO_MICROTUBULE_C | 43 | 2.3322495613 | 0 | 0 |
| GO_REGULATION_OF | 87 | 2.3645791588 | 0 | 0 |

### Broad GSEA 1.0.0

The species: **"human"**
The database for the enrichement analysis: **"go"**
Number of repeats for calculating p-values: **1000**
A list of genes with their scores (usually output from scanpy scoring for example): **Marker score (Cluster 2 vs. rest)** (from step 12: Marker Genes Identification 1.0.0)

| gset_name | gset_size | nes | p_val | fdr |
|---|---|---|---|---|
| GO_CALCIUM_DEPEN | 12 | 3.4634105437 | 0 | 0 |
| GO_CELL_PROJECTI( | 139 | 2.3612219797 | 0 | 0 |
| GO_GLUCOSE_METAI | 21 | 3.4714676181 | 0 | 0 |
| GO_OXIDATION_REDU | 78 | 2.8874490122 | 0 | 0 |
| GO_NEGATIVE_REGU | 14 | 3.5490548993 | 0 | 0 |
| GO_ACTIN_FILAMENT | 50 | 2.7609463215 | 0 | 0 |
| GO_PURINE_CONTAII | 35 | 3.6790812319 | 0 | 0 |
| GO_STRUCTURAL_C( | 18 | 3.8285672551 | 0 | 0 |
| GO_CARBOHYDRATE_ | 54 | 2.6840172988 | 0 | 0 |

### Broad GSEA 1.0.0

The species: **"human"**
The database for the enrichement analysis: **"go"**
Number of repeats for calculating p-values: **1000**
A list of genes with their scores (usually output from scanpy scoring for example): **Marker score (Cluster 3 vs. rest)** (from step 12: Marker Genes Identification 1.0.0)

| gset_name | gset_size | nes | p_val | fdr |
|---|---|---|---|---|
| GO_NEGATIVE_REGU | 88 | 2.5290871941 | 0 | 0 |
| GO_ADENYL_NUCLE( | 110 | 2.4677889845 | 0 | 0 |
| GO_POSITIVE_REGUL | 17 | 4.0281776068 | 0 | 0 |
| GO_ORGANELLE_FIS! | 46 | 4.3648616611 | 0 | 0 |
| GO_CHROMATIN_OR( | 34 | 3.3637933508 | 0 | 0 |
| GO_DNA_PACKAGIN( | 18 | 3.6002319314 | 0 | 0 |
| GO_SPINDLE_POLE | 16 | 4.015371666 | 0 | 0 |
| GO_CHROMATIN | 38 | 3.2656739887 | 0 | 0 |
| GO_POSITIVE_REGUL | 83 | 2.5615805609 | 0 | 0 |

### Pseudotime construction 1.0.0

Number of neighbors to calculate: **20**
Method for computing connectivities: **"gauss"**
The input assay to use: **Log transformed assay** (from step 5: Log transformation 1.0.0)

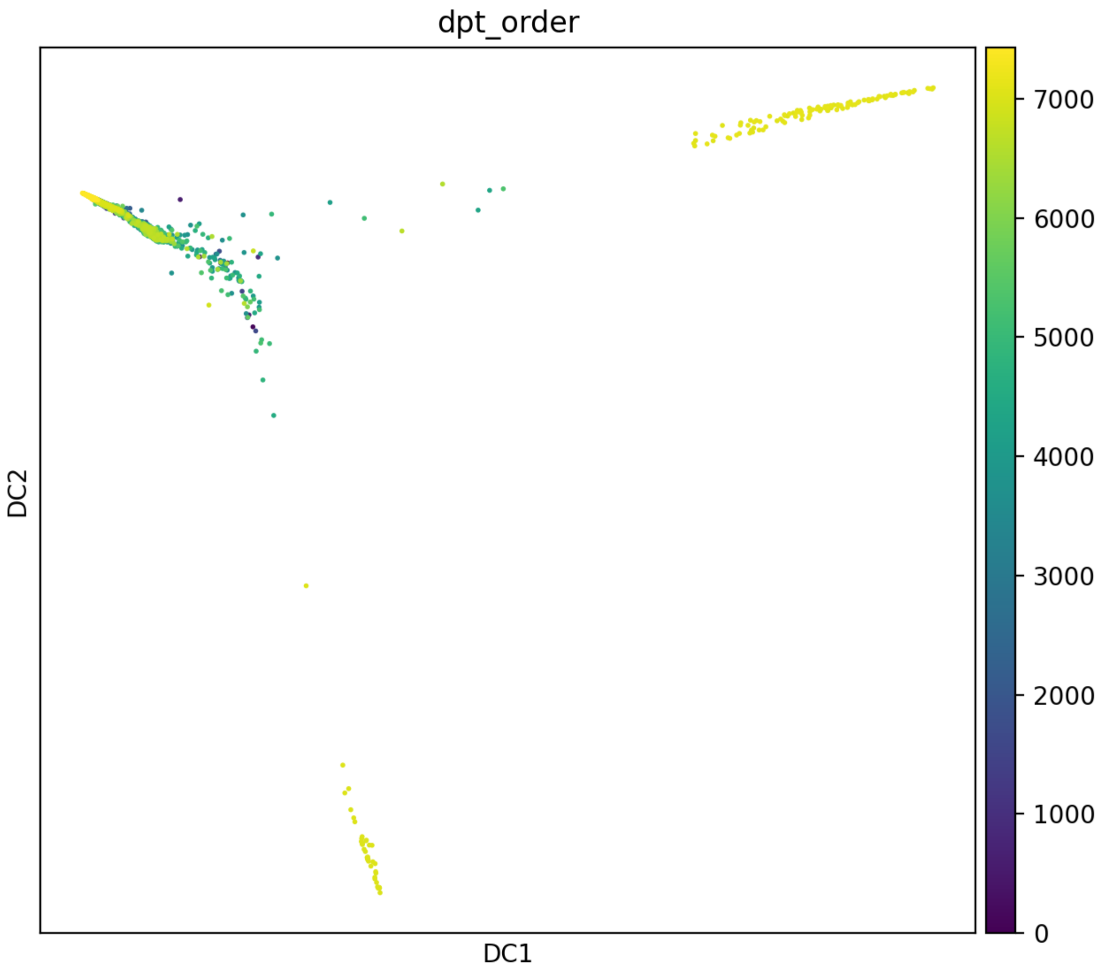


Cell order

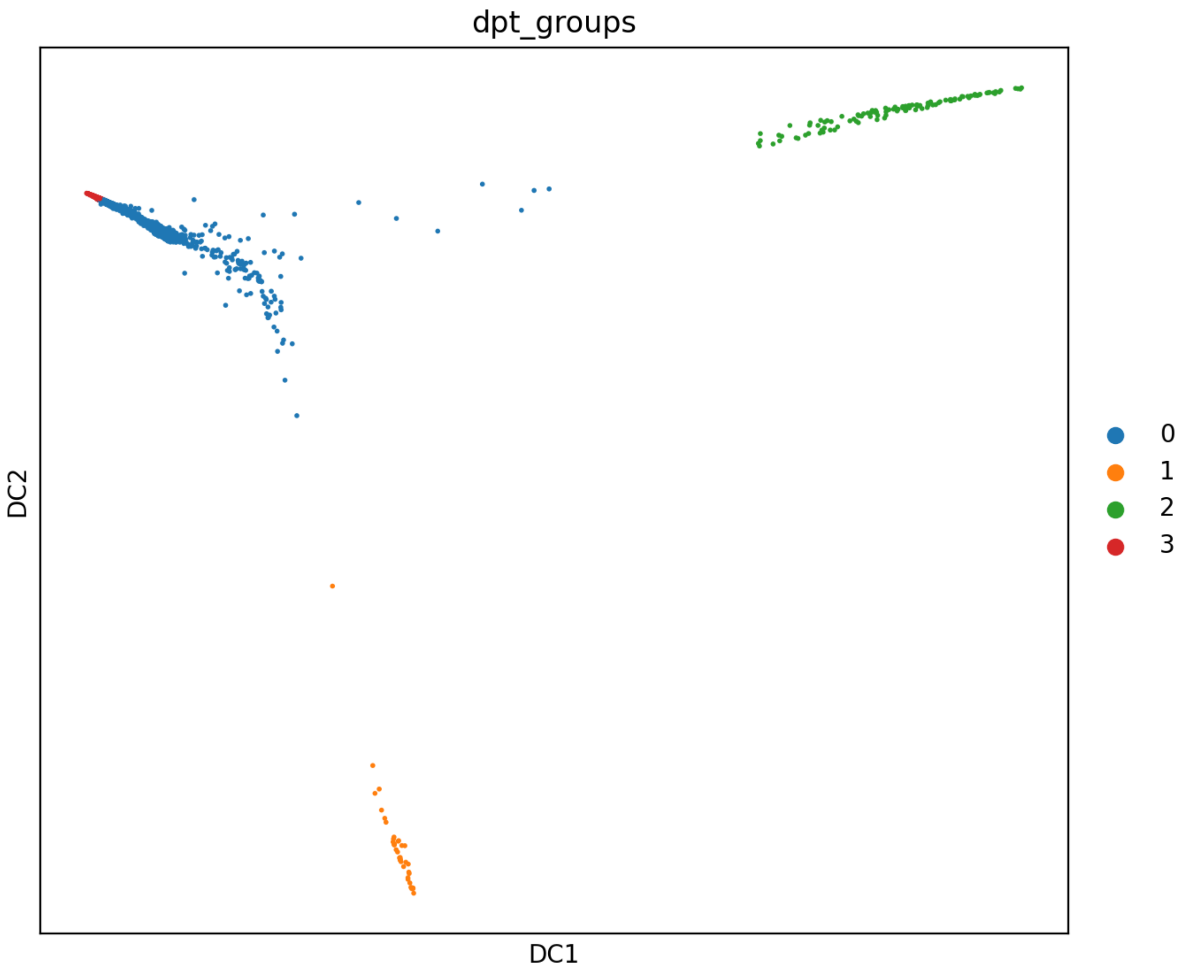


Cell groups

Use the browser back button to return to the project steps.
