## Supplementary File 3 for "GranatumX: A Community-engaging, Modularized, and Flexible Webtool for Single-cell Data Analysis"

### Manuscript pipeline

#### Analysis Report

This report is generated by GranatumX

Please cite: Zhu, Xun, et al. "GranatumX: A community engaging and flexible software environment for single-cell analysis." bioRxiv (2018): 385591.

### Upload Files 1.0.0

Assay to upload: **TM_droplet.csv.zip (107.64 MB)**
Sample metadata file to upload: **annotations_droplet.csv (8.92 MB)**
File format: **"zip"**
Convert gene IDs: **false**
Species: **"mouse"**
Convert gene IDs into (HGNC symbol is recommended): **"symbol"**
Add extra info (from BioMart) into gene metadata: **true**
Enter your email address to get notified of any errors encountered in the pipeline: **false**

The assay has **23433** genes (with inferred ID type: Symbol) and **54865** samples.

The first few rows and columns:

```
0.0, 0.0, 0.0, 0.0, 0.0, 0.0, 0.0, 0.0, 0.0, 0.0
3.0, 1.0, 3.0, 0.0, 1.0, 1.0, 1.0, 0.0, 0.0, 1.0
0.0, 1.0, 0.0, 0.0, 0.0, 1.0, 1.0, 1.0, 0.0, 1.0
0.0, 0.0, 0.0, 0.0, 0.0, 0.0, 0.0, 0.0, 0.0, 0.0
2.0, 0.0, 0.0, 0.0, 0.0, 1.0, 0.0, 0.0, 0.0, 0.0
1.0, 0.0, 0.0, 0.0, 2.0, 1.0, 2.0, 0.0, 0.0, 0.0
0.0, 3.0, 0.0, 2.0, 0.0, 0.0, 0.0, 1.0, 0.0, 0.0
0.0, 0.0, 0.0, 0.0, 0.0, 0.0, 0.0, 0.0, 0.0, 0.0
0.0, 0.0, 0.0, 0.0, 0.0, 0.0, 0.0, 0.0, 0.0, 0.0
2.0, 0.0, 1.0, 2.0, 2.0, 0.0, 0.0, 3.0, 0.0, 2.0
```

### DeepImpute 2.0.0

Random seed: **12345**
Use automatic gene imputation limit: **true**
Gene rank limit: **2000**
Cell subset: **1**
Assay: **[A]TM_droplet.csv.zip** (from step 1: Upload Files 1.0.0)

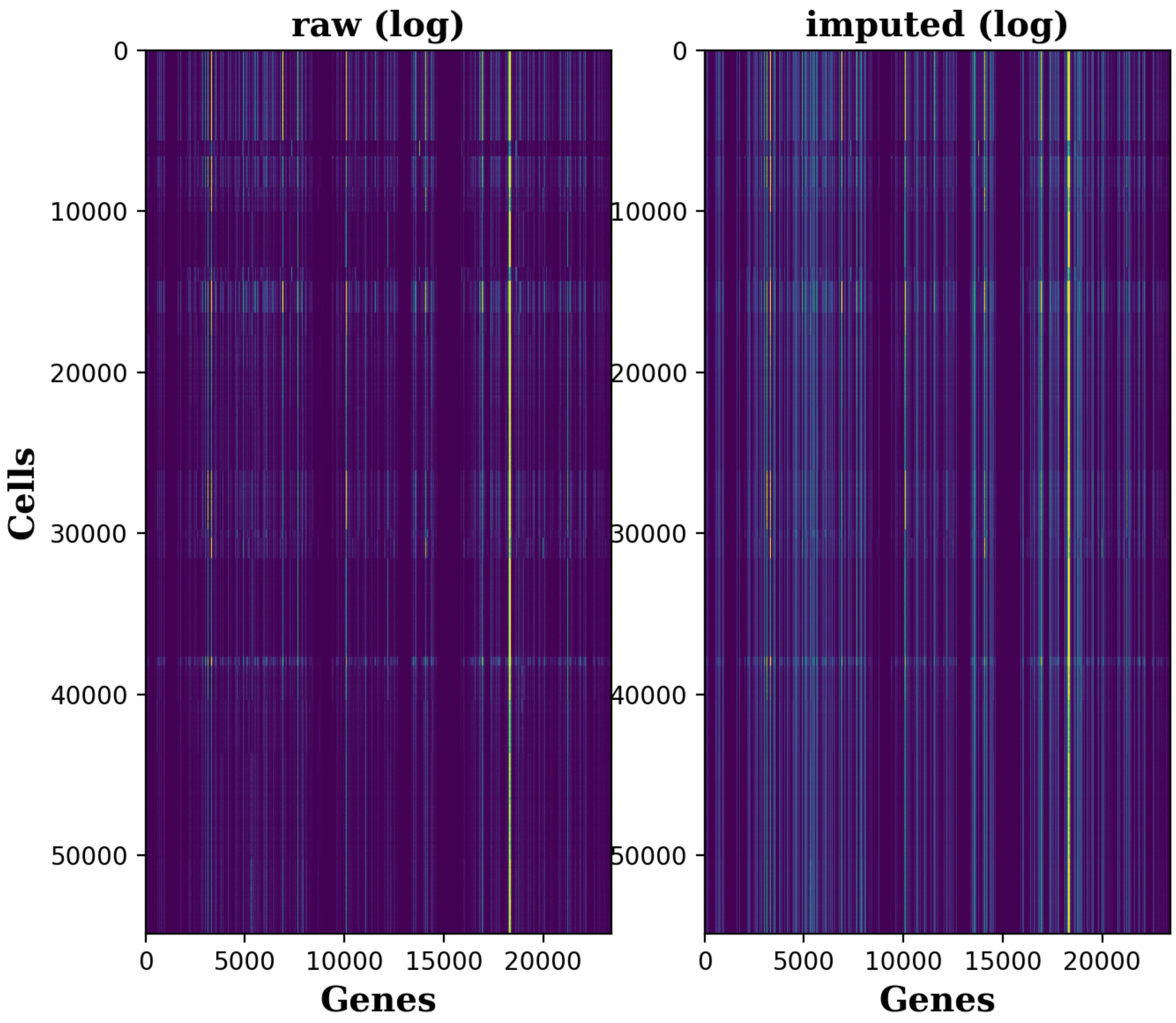


Heatmaps

- Data frame number of rows: **54865**
- Data frame number of columns: **23433**
- Number of imputed genes: **3584**

- Percentage of dropout entries *before* imputation: **91.82%**
- Percentage of dropout entries *after* imputation: **81.17%**
- Accuracy (correlation) on masked data: **0.95**

### Cell Normalization 1.0.0

Log transform in the boxplots: **true**
Normalization method: **"quantile"**
Number of cells to plot in the bar-plot: **40**
Assay: **Imputed assay** (from step 2: DeepImpute 2.0.0)

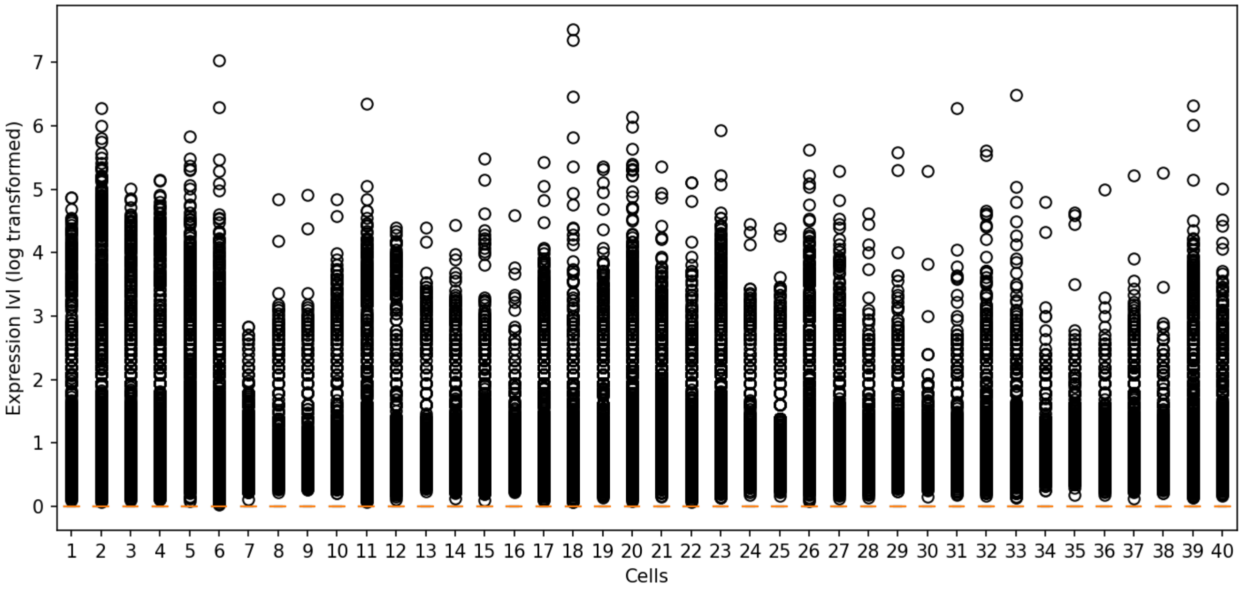


Before normalization: Each bar in the box plot represents one cell.

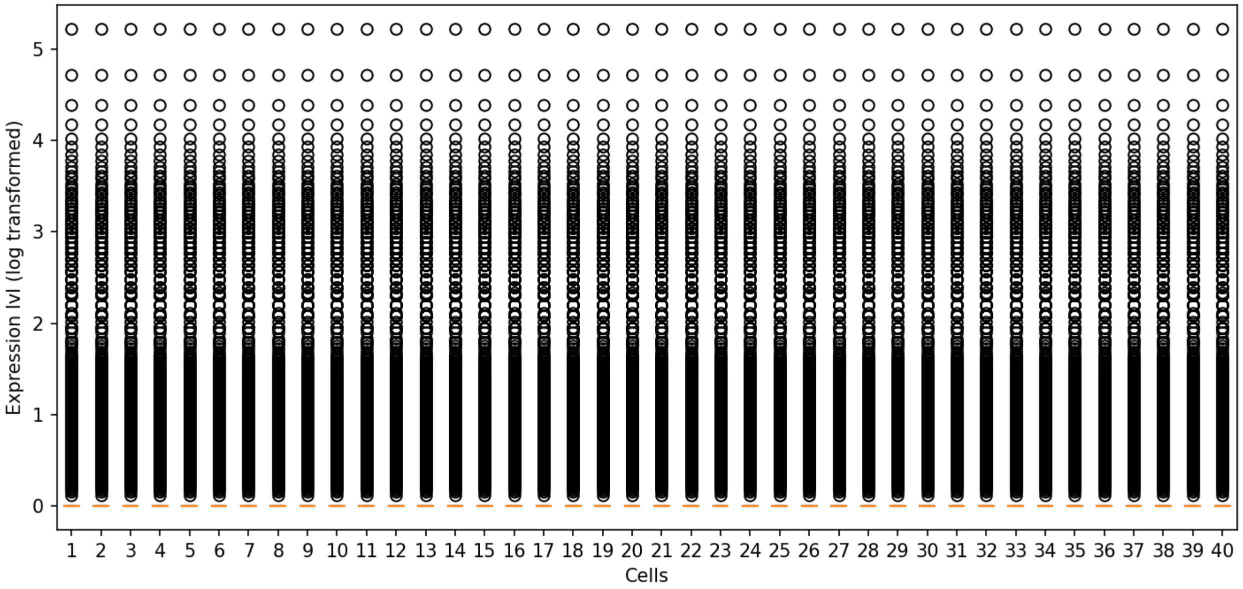


After normalization: Each bar in the box plot represents one cell.

### Scanpy Gene Filtering 1.0.0

The gene has to be expressed in at least ___ cells: **3**
The average expression level of the gene has to be greater than: **1**
The average expression level of the gene has to be less than: **999999**
The dispersion of the gene has to be greater than: **0.5**
The dispersion of the gene has to be less than: **999999**
Assay: **Normalized assay** (from step 3: Cell Normalization 1.0.0)

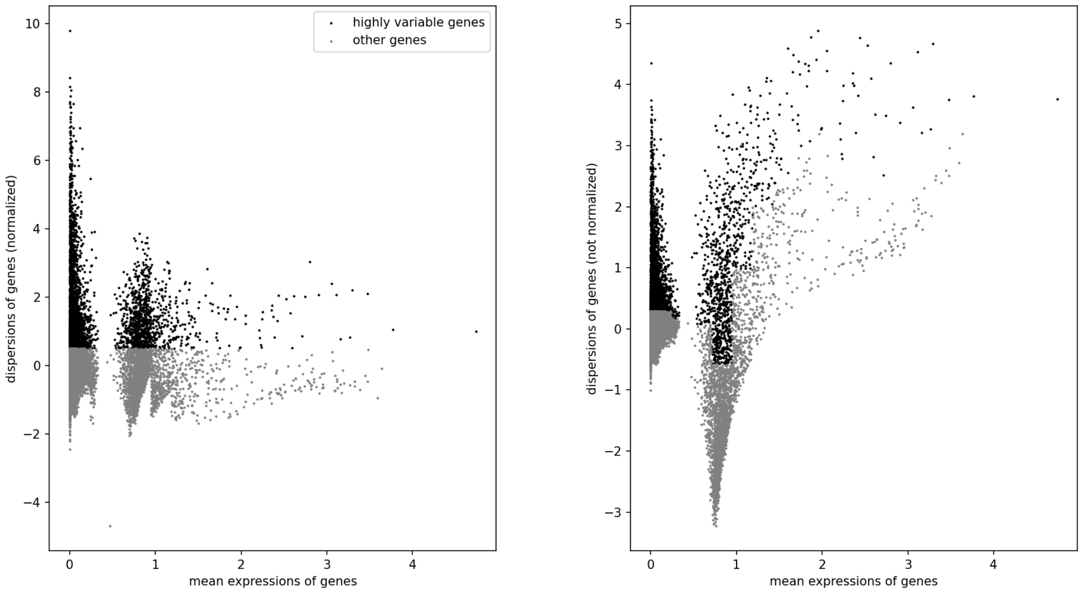


Each dot represent a gene. The gray dots are the removed genes. The x-axis is log-transformed.

Number of genes before filtering: **23433**

Number of genes after filtering: **3531**

### Log transformation 1.0.0

The base used for the log function: **2**

The pseudo counts added before log transformation (to avoid getting log(0)): **1**

Assay including matrix and geneIds: **Filtered Assay** (from step 4: Scanpy Gene Filtering 1.0.0)

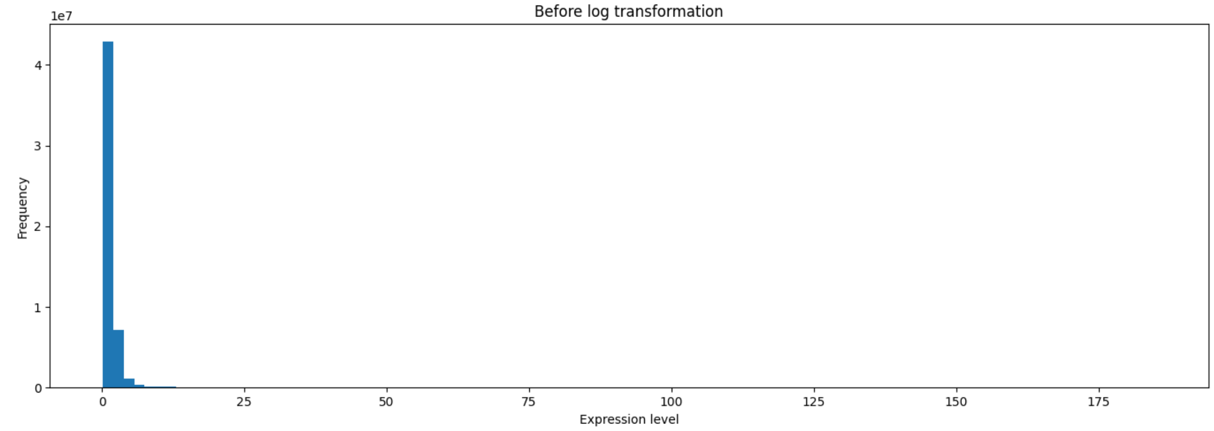


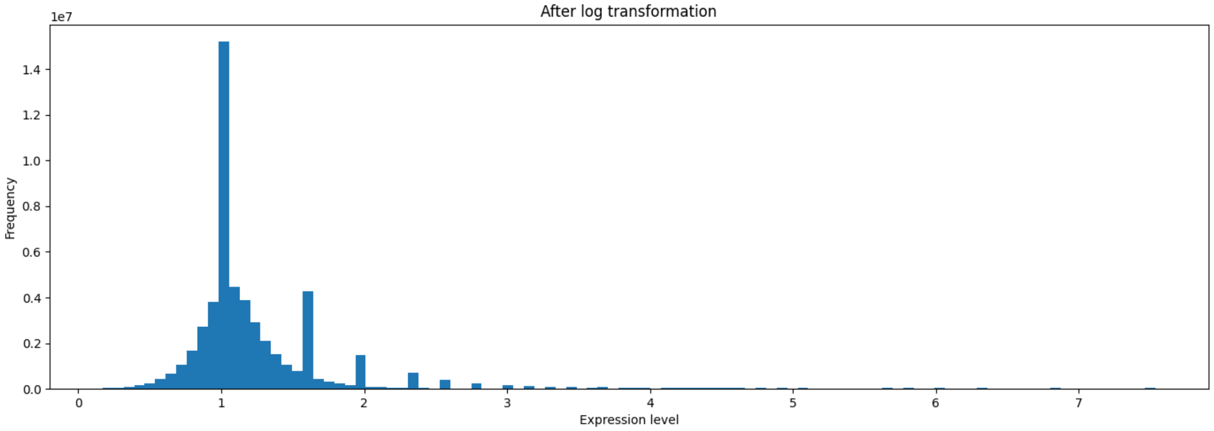


The distribution of expression level before and after log transformation. Only the values greater than the 5 percentile (usually zero in single-cell data) and lower than 95 percentile are considered.

### Principal Component Analysis 1.0.0

Number of top components to calculate: **2**

Assay: **Log transformed assay** (from step 5: Log transformation 1.0.0)

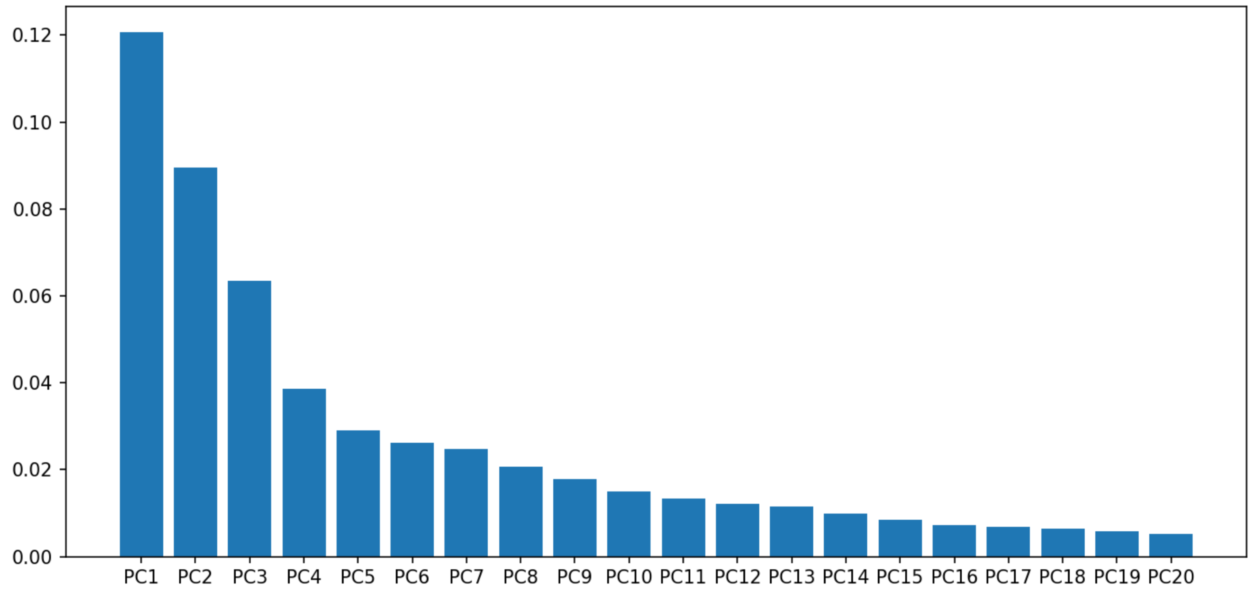


Explained variance (ratio) by each Principal Component (PC)

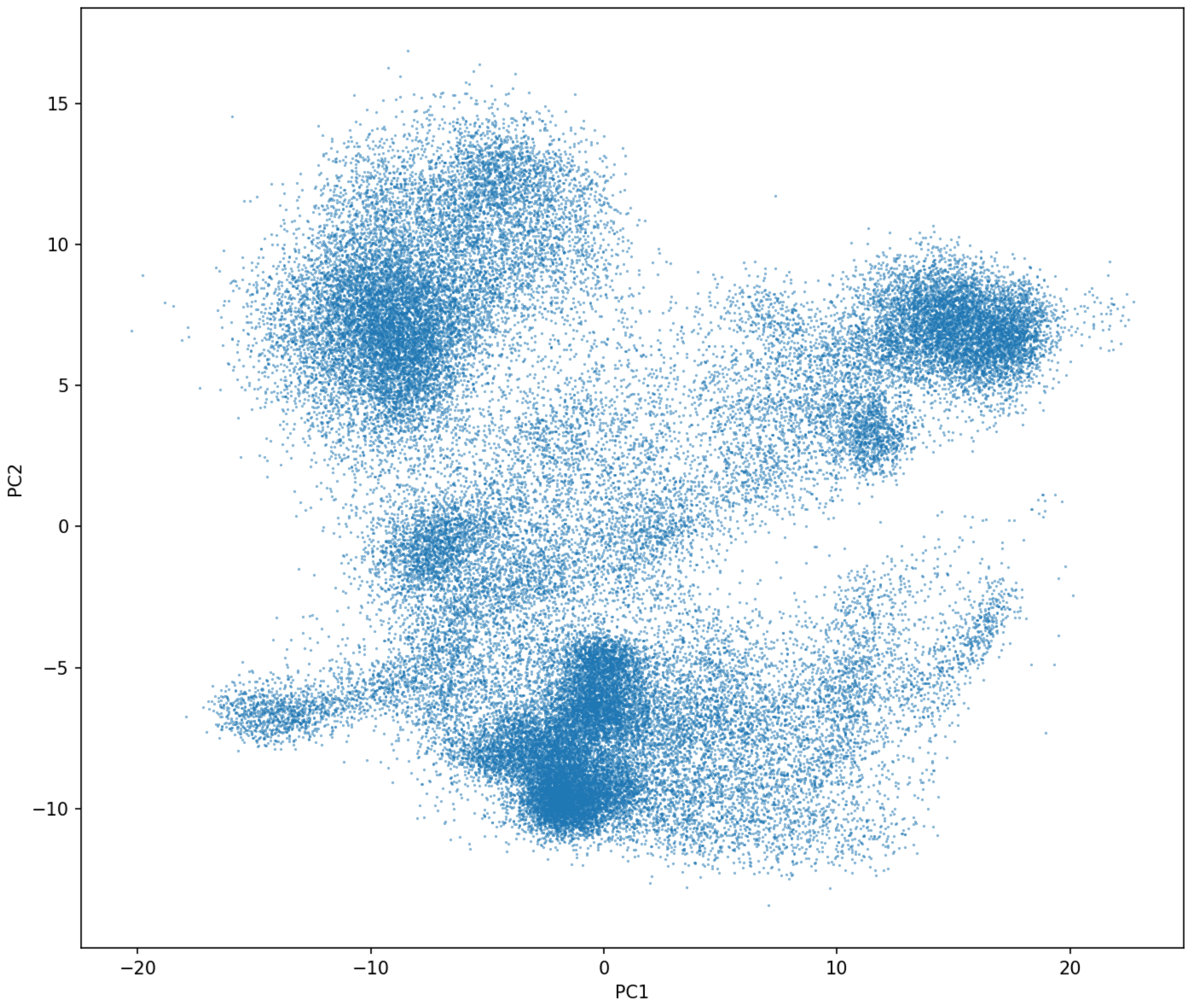


PC1 vs. PC2

### UMAP 1.0.0

Number of neighbors (n_neighbors, an integer): **15**
Minimum distance (min_dist, a real number ranges from 0 to 1): **0.1**
Metric (metric): **"euclidean"**
Assay: **Log transformed assay** (from step 5: Log transformation 1.0.0)

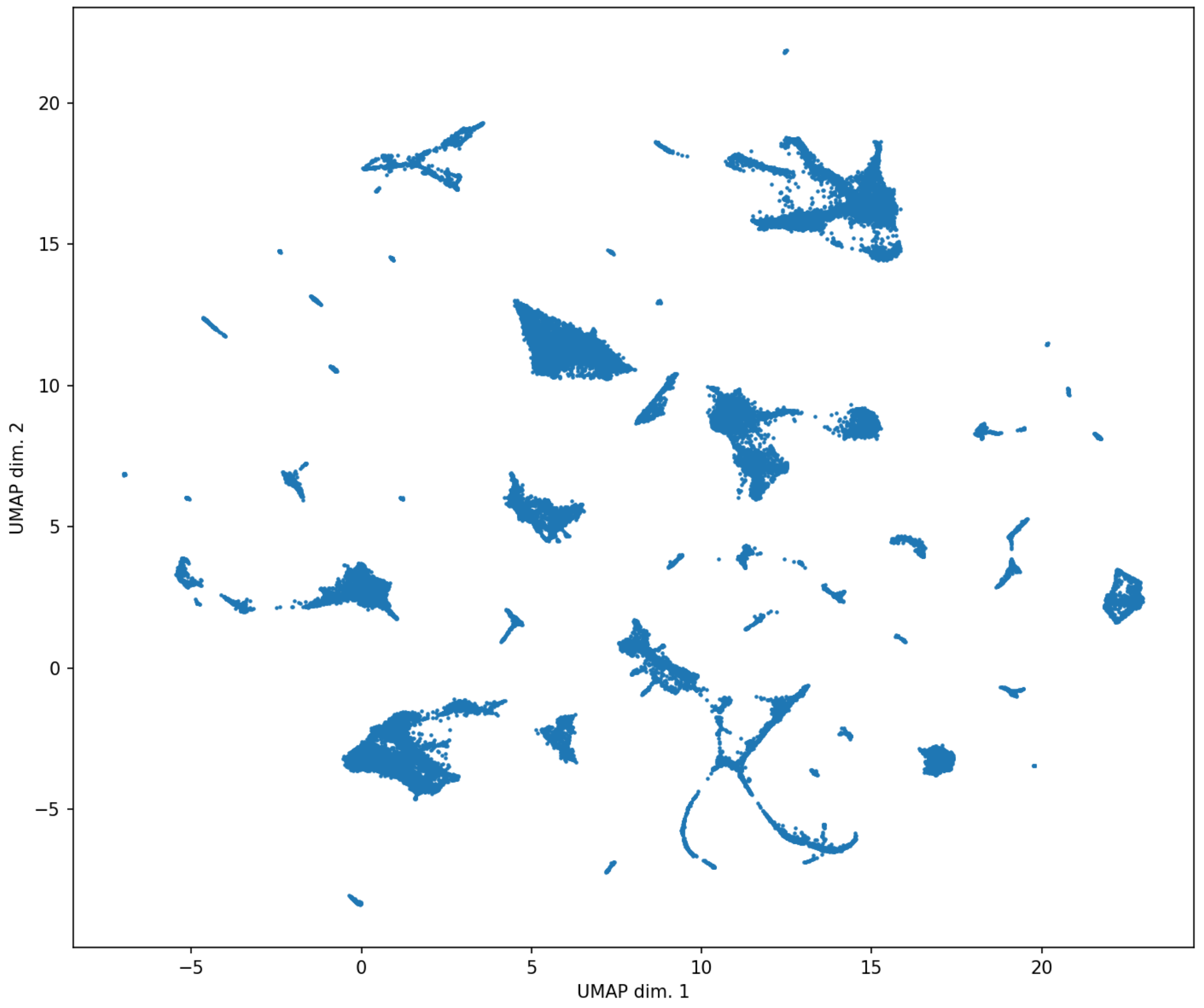


UMAP plot: each dot represents a cell

### ScanpyClustering 1.0.0

Random seed: **13513**

Assay including matrix and geneIds: **Log transformed assay** (from step 5: Log transformation 1.0.0)

Cell coordinates for visualization: **UMAP coordinates** (from step 7: UMAP 1.0.0)

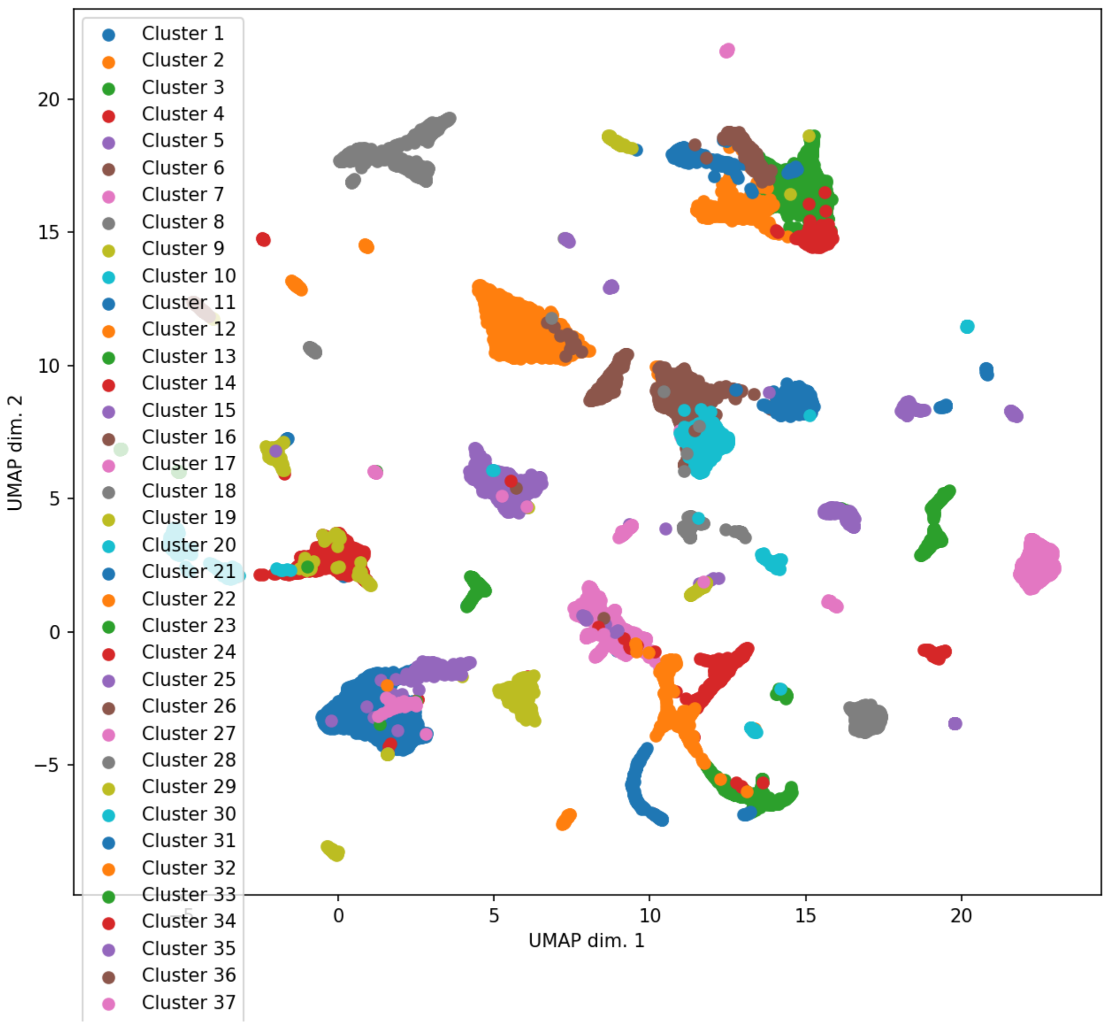


Scatter-plot using imported cell coordinates. Each dot represents a cell. The colors indicate the indentified cell clusters.

### Sample Coloring 1.0.0

The type of the sample metadata: **"categorical"**
Bounding box factor times standard deviation (value should be > 0.0; 0.0 will be the convex hull): **0.1**
The fontsize of the labels: **9**
Values to use as colors: **Cluster assignment** (from step 8: ScanpyClustering 1.0.0)
Visualization data to plot: **UMAP coordinates** (from step 7: UMAP 1.0.0)

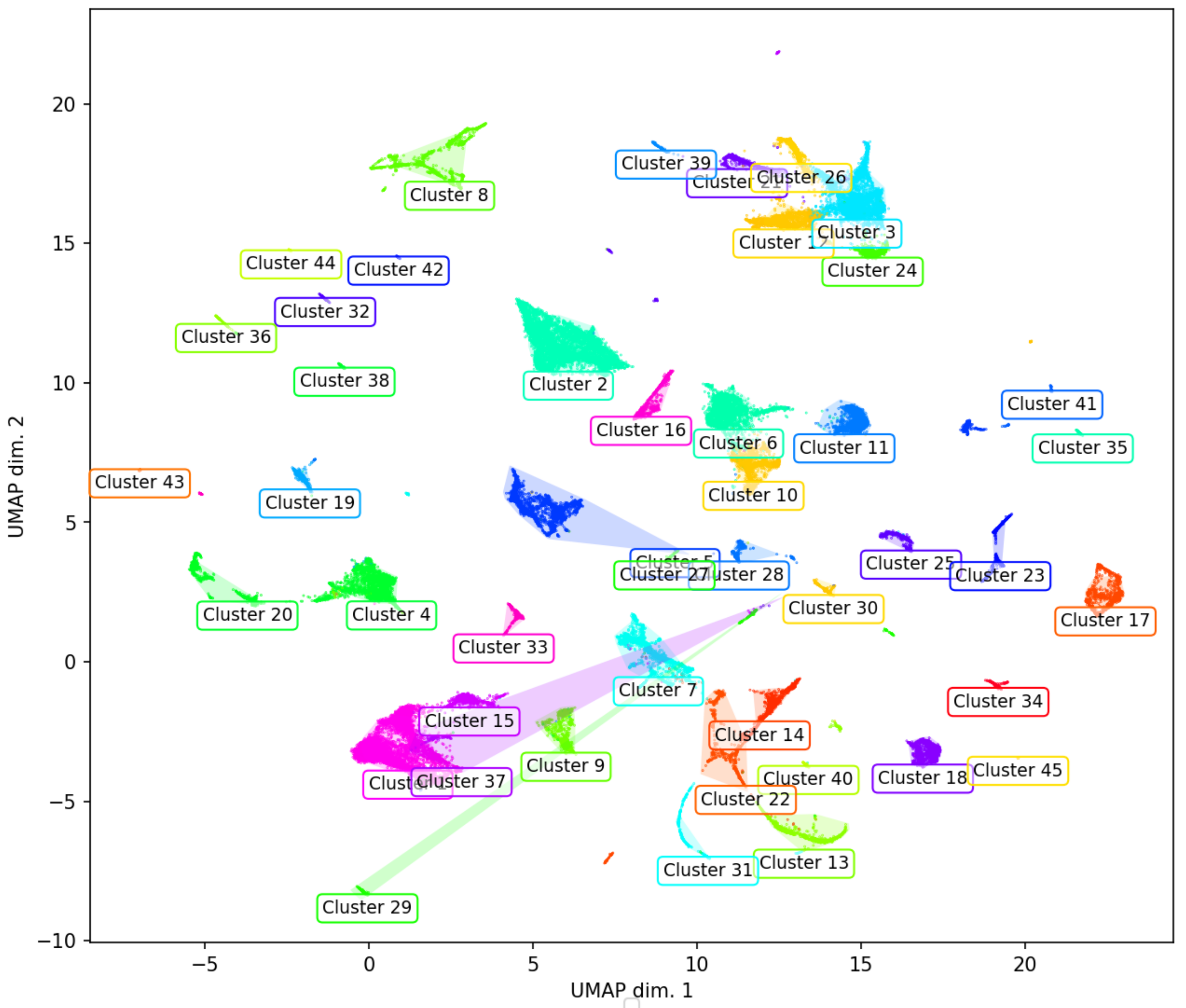


Scatter-plot

### Sample Coloring 1.0.0

The type of the sample metadata: **"categorical"**
Bounding box factor times standard deviation (value should be > 0.0; 0.0 will be the convex hull): **0.1**
The fontsize of the labels: **9**
Values to use as colors: **[M]tissue** (from step 1: Upload Files 1.0.0)
Visualization data to plot: **UMAP coordinates** (from step 7: UMAP 1.0.0)

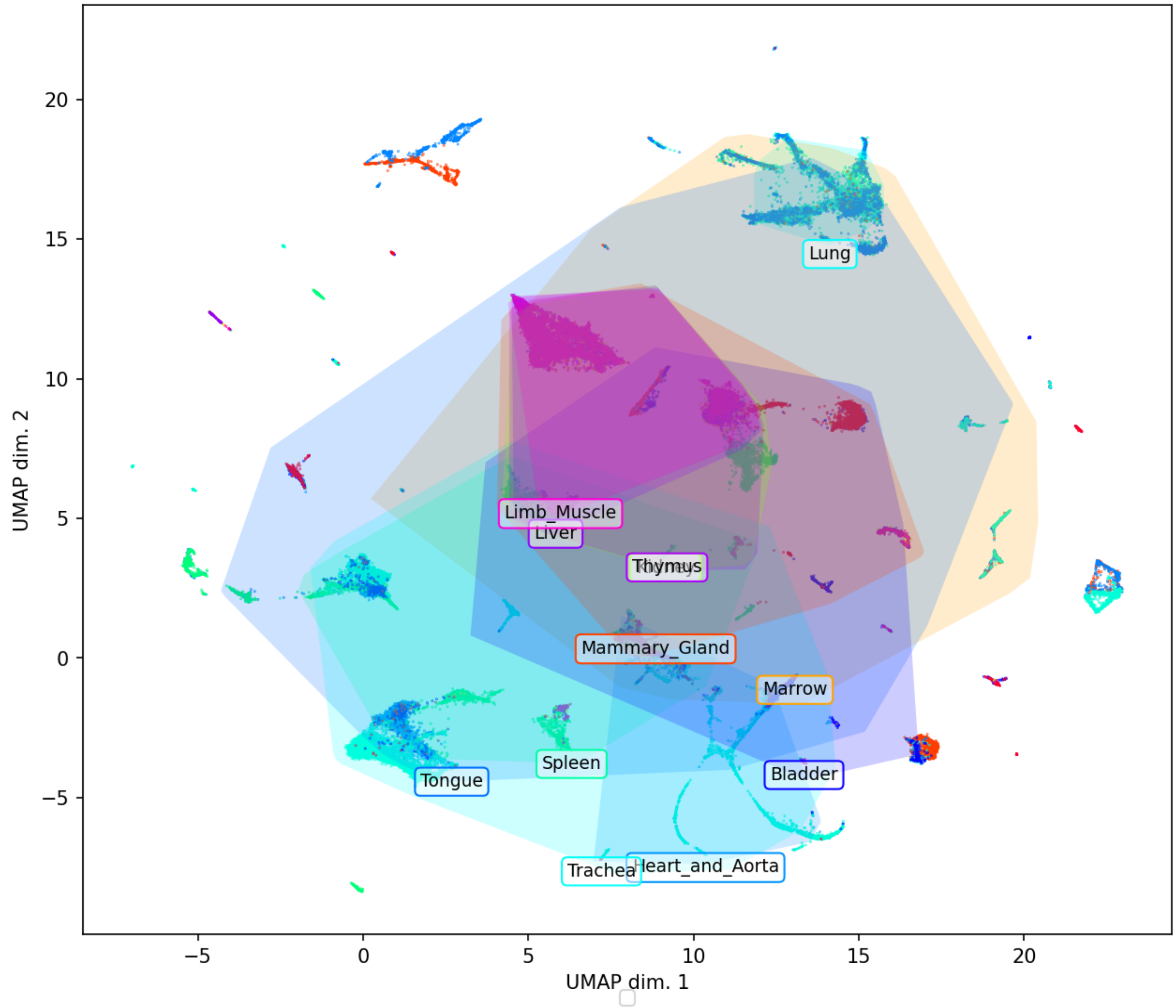


Scatter-plot

### Marker Genes Identification 1.0.0

Assay including matrix and geneIds: **Log transformed assay** (from step 5: Log transformation 1.0.0)
Group vector: **[M]tissue** (from step 1: Upload Files 1.0.0)

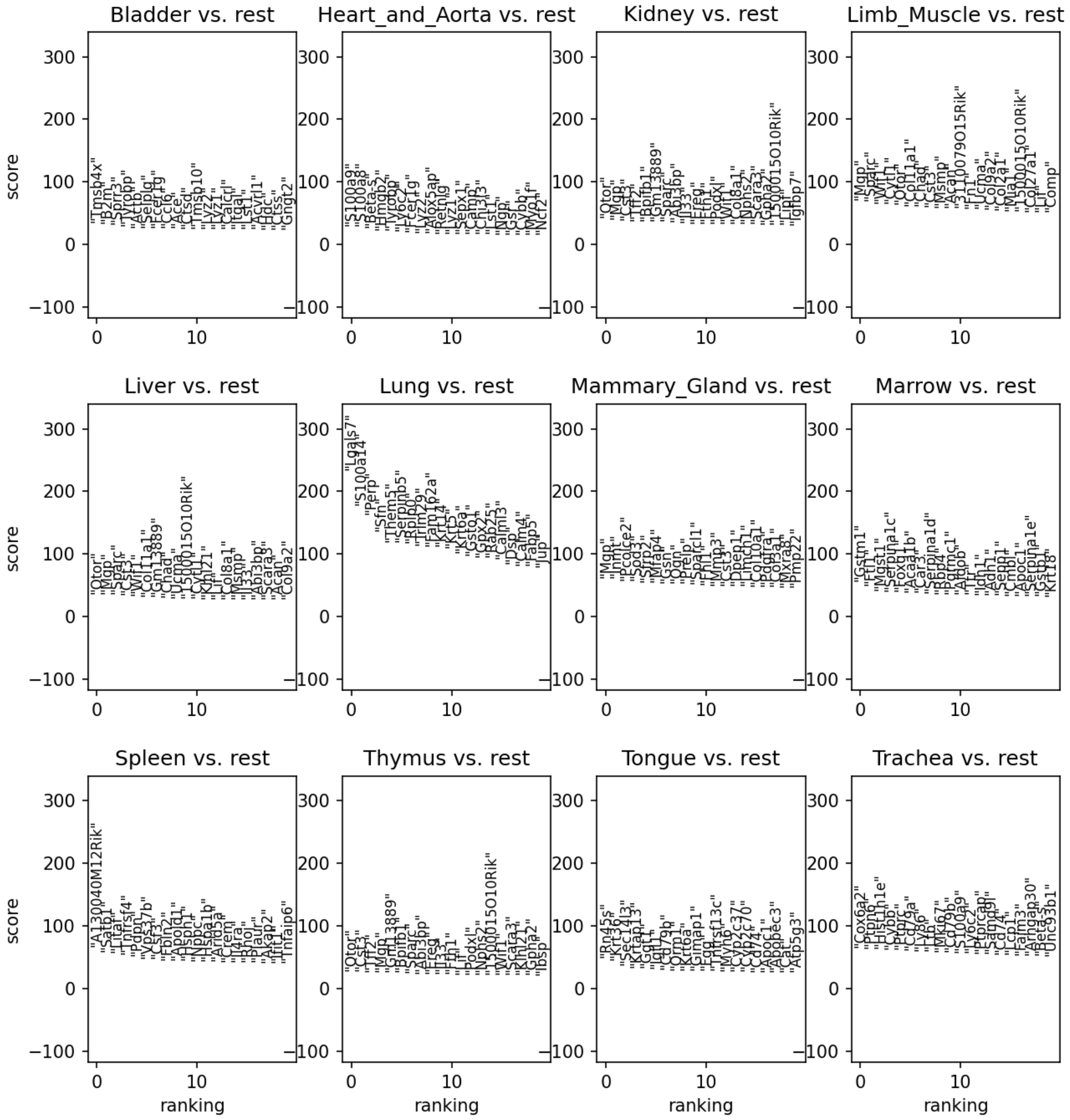


One-vs-rest marker genes

### Marker Genes Identification 1.0.0

Assay including matrix and geneIds: **Log transformed assay** (from step 5: Log transformation 1.0.0)
Group vector: **Cluster assignment** (from step 8: ScanpyClustering 1.0.0)

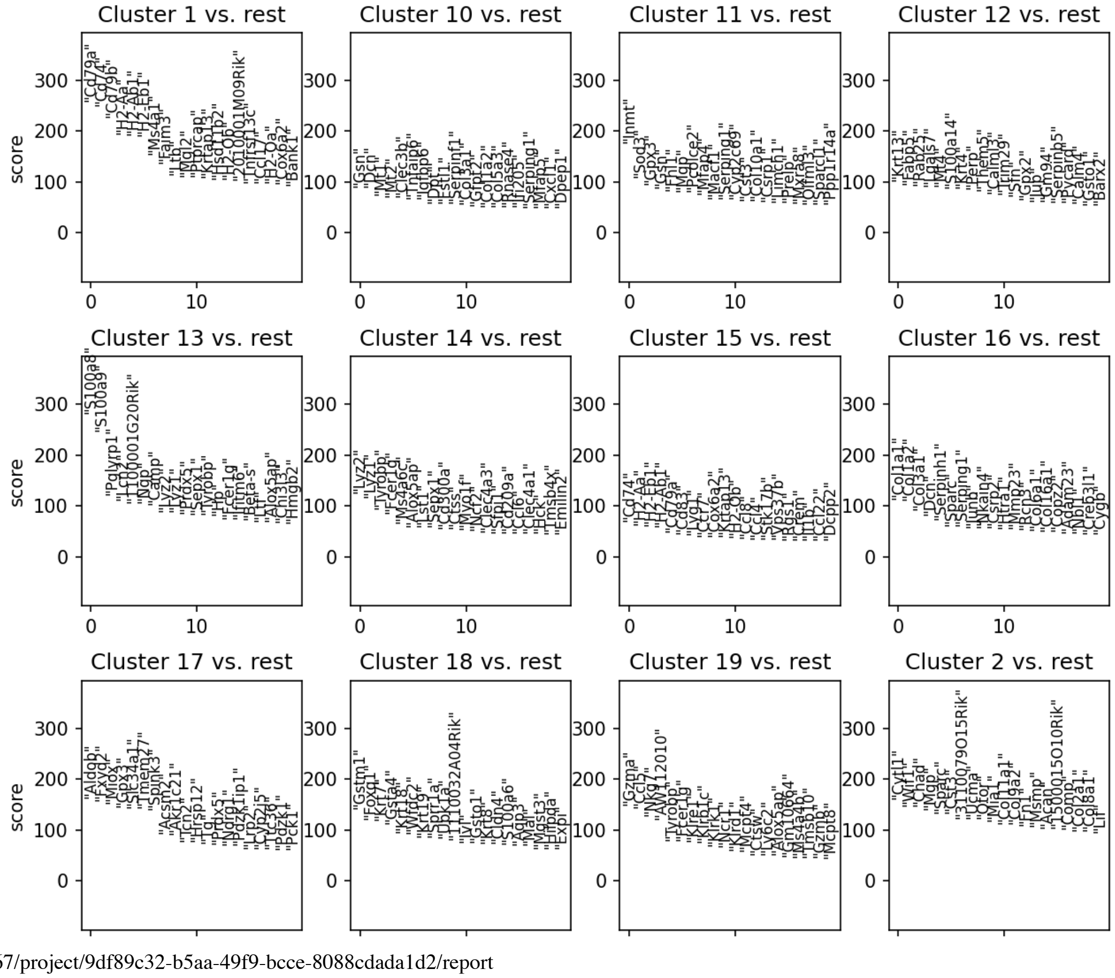

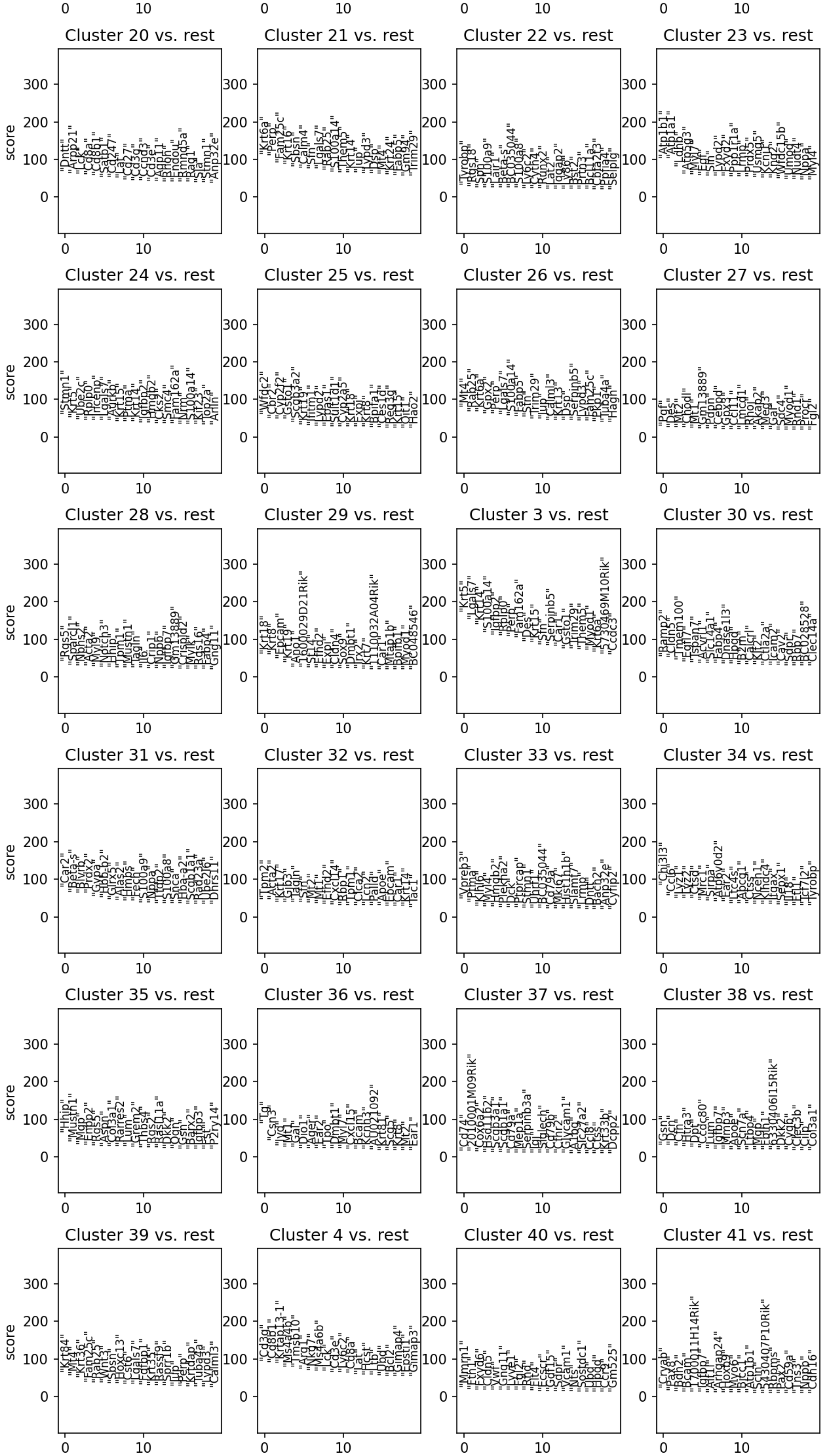
Cluster 20 vs. rest
Cluster 21 vs. rest
Cluster 22 vs. rest
Cluster 23 vs. rest
Cluster 24 vs. rest
Cluster 25 vs. rest
Cluster 26 vs. rest
Cluster 27 vs. rest
Cluster 28 vs. rest
Cluster 29 vs. rest
Cluster 3 vs. rest
Cluster 30 vs. rest
Cluster 31 vs. rest
Cluster 32 vs. rest
Cluster 33 vs. rest
Cluster 34 vs. rest
Cluster 35 vs. rest
Cluster 36 vs. rest
Cluster 37 vs. rest
Cluster 38 vs. rest
Cluster 39 vs. rest
Cluster 4 vs. rest
Cluster 40 vs. rest
Cluster 41 vs. rest
score
0
100
200
300
10

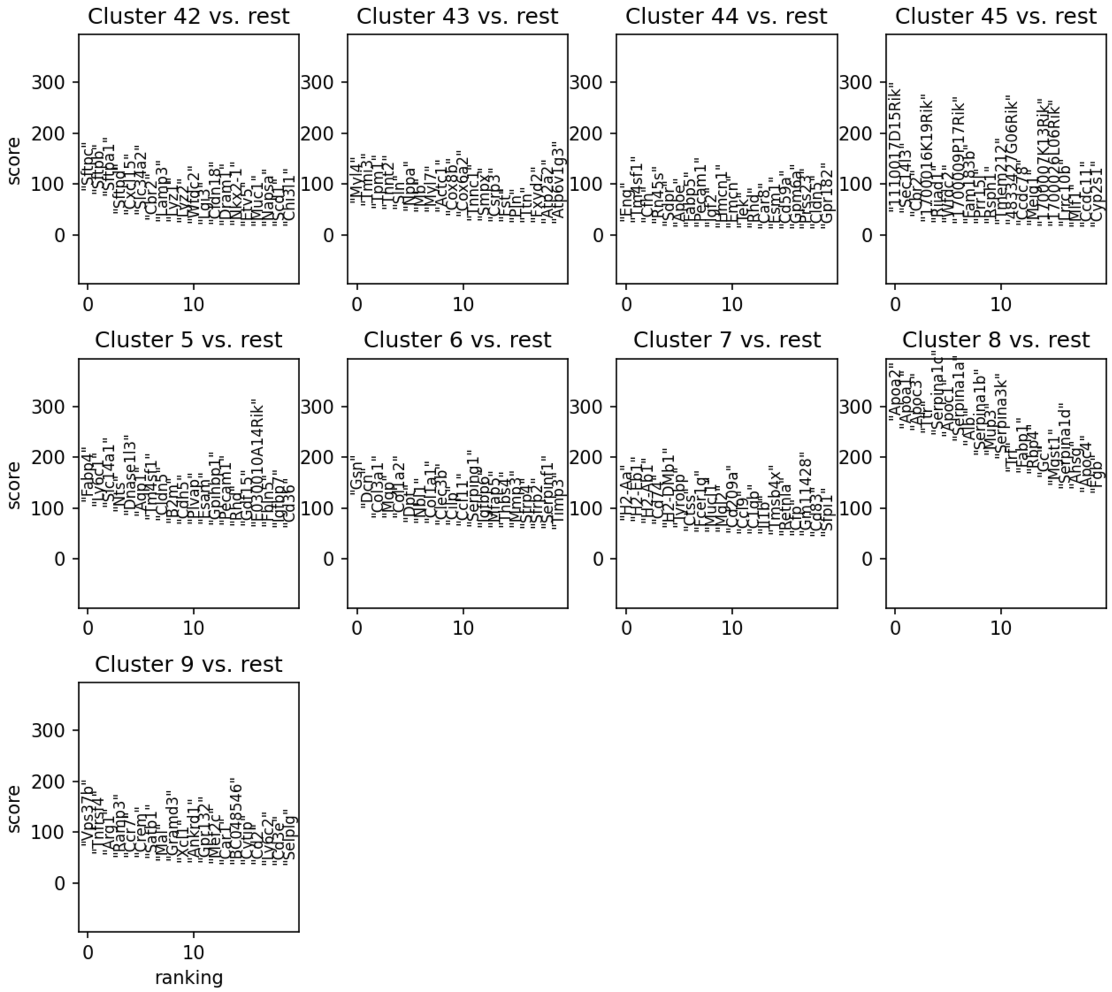


One-vs-rest marker genes

Use the browser back button to return to the project steps.
